# Both YAP1-MAML2 and constitutively active YAP1 drive the formation of tumors that resemble NF2-mutant meningiomas in mice

**DOI:** 10.1101/2022.05.02.490337

**Authors:** Frank Szulzewsky, Sonali Arora, Aleena Arakaki, Philipp Sievers, Damian A. Almiron Bonnin, Patrick J. Paddison, Felix Sahm, Patrick J Cimino, Taranjit S Gujral, Eric C Holland

## Abstract

YAP1 is a transcriptional co-activator regulated by the Hippo Signaling Pathway, including NF2. Meningiomas are the most common primary brain tumors, a large percentage exhibit heterozygous loss of chromosome 22 (harboring the *NF2* gene) and functional inactivation of the remaining *NF2* copy, implicating oncogenic YAP activity in these tumors. Recently, fusions between YAP1 and MAML2 have been identified in a subset of pediatric *NF2*-wild type meningiomas. Here, we show that human *YAP1-MAML2*-positive meningiomas resemble *NF2*-mutant meningiomas by global and YAP-related gene expression signatures. We then show that expression of *YAP1-MAML2* in mice induces tumors that resemble human YAP1 fusion-positive and *NF2*-mutant meningiomas by gene expression. We demonstrate that YAP1-MAML2 primarily functions by exerting TEAD-dependent YAP activity that is resistant to Hippo signaling. Treatment with YAP-TEAD inhibitors is sufficient to inhibit the viability of *YAP1-MAML2*-driven mouse tumors *ex vivo*. Finally, we show that expression of constitutively active YAP1 (S127/397A-YAP1) is sufficient to induce similar tumors suggesting that the YAP component of the gene fusion is the critical driver of these tumors. In summary, our results implicate YAP1-MAML2 as a causal oncogenic driver and highlight TEAD-dependent YAP activity as an oncogenic driver in *YAP1-MAML2*-fusion meningioma as well as NF2-mutant meningioma in general.

## Introduction

Meningiomas are the most common primary brain tumors in adults, accounting for 36.4 percent of all cases, whereas pediatric cases are rare (Ostrom et al. 2020). Around half of these tumors exhibit functional loss of the tumor suppressor *NF2*. Of the remaining *NF2*-wildtype meningiomas, half harbor mutations in *TRAF7, KLF4, AKT1*, or *SMO*. The vast majority of meningiomas are benign, but higher-grade tumors and recurrences occur in all molecular subgroups (Sahm et al. 2017). Recent genomewide DNA methylation studies identified clinically relevant meningioma classes correlating typical mutational, cytogenetic, and gene expression patterns with clinical outcomes (Sahm et al. 2017; Nassiri et al. 2021). Mutations in *TRAF7, KLF4, AKT1*, and *SMO* are restricted to a specific benign subtype, whereas *NF2*-mutant tumors are enriched in all other subtypes. While benign *NF2*-mutant tumors show virtually no copy number alterations in addition to the heterozygous loss of chromosome 22, mutational burden and additional chromosomal aberrations are more frequent in malignant subtypes and higher-grade tumors. Historically, losses or functional inactivation of known tumor suppressor genes (such as *CDKN2A/B*) have been reported in all grades but were more frequent in higher-grade tumors (Barresi et al. 2021). Homozygous deletion of *CDKN2A/B* has also been associated with clinical recurrence of meningioma (Sievers et al. 2020b) and is now considered a molecular criterion sufficient for the diagnosis of Anaplastic (Malignant) Meningioma, WHO Grade 3 (Louis et al. 2021).

YAP1 and its paralogue TAZ (encoded by *WWTR1*) are transcriptional co-activators and potent drivers of cell growth that function through the interaction with several different transcription factors, most prominently TEAD1-4 (Szulzewsky et al. 2021). The activity of YAP1 is regulated by the Hippo signaling pathway, a cascade of serine/threonine kinases that ultimately phosphorylate YAP1 at several serine residues, resulting in the inhibition of YAP activity. Elevated and nuclear YAP1 staining has been observed in several cancers and inactivating mutations in upstream Hippo Pathway tumor suppressors (such as *NF2, FAT1-4*, or *LATS1/2*) occur in a multitude of cancers. *NF2*/Merlin is a potent positive regulator of the Hippo pathway (and therefore an inhibitor of YAP1) and inactivating mutations in the *NF2* gene – in addition to the high prevalence in meningioma – are also frequently found in schwannoma, ependymoma, and malignant mesothelioma. The high prevalence of heterozygous deletions of chromosome 22 (harboring the *NF2* gene) and additional functional inactivation of the remaining *NF2* copy in meningiomas causally implicates de-regulated and oncogenic YAP activity in the pathobiology of these tumors. Strikingly, the development of hepatocellular carcinomas in *Nf2* knockout mice was dependent on the presence of functional YAP1 (Zhang et al. 2010), further linking NF2 inactivation to oncogenic YAP1 signaling.

Genetically engineered mouse models (GEMMs) of human cancers are valuable tools for pre-clinical drug testing and for studying the underlying oncogenic drivers and molecular pathways in these tumors. While the majority of meningiomas are benign, a subset are unresponsive to therapy, recur even after multiple surgeries, and are ultimately lethal. Hence, GEMMs of meningiomas are needed to develop better treatments for these tumors. Unfortunately, the generation of meningioma GEMMs has been hampered by the lack of strong oncogenic drivers that can be used for modeling this disease and the benign nature and slow growth of a majority of these tumors (Kalamarides et al. 2002; Peyre et al. 2013), rendering these models challenging for pre-clinical research.

Recently, YAP1 gene fusion events retaining the N-terminal domains of YAP1, including the TEAD-interacting domain, have been identified in a subset of pediatric *NF2*-wild type meningiomas (Sievers et al. 2020a). *YAP1-MAML2* was identified in 7 out of 9 cases, while two additional YAP1 fusions (*YAP1-PYGO1* and *YAP1-LMO1*) were each identified in one case. *YAP1-MAML2* has also been identified in several other cancers, such as Poroma/Porocarcinoma, retiform and composite hemangioendothelioma; Head and Neck, Nasopharyngeal, and Ovarian Carcinoma (Sekine et al. 2019; Antonescu et al. 2020; Szulzewsky et al. 2021). Additional YAP1 gene fusions have been identified in several cancer subtypes, including Supratentorial Ependymoma and Epithelioid Hemangioendothelioma (Szulzewsky et al. 2021). YAP1 fusion proteins exhibit conserved structural and functional features, most importantly their ability to exert TEAD-dependent YAP activity that is resistant to inhibitory Hippo signaling and we and others have previously shown that several of these YAP1 gene fusions are oncogenic when expressed in mice (Pajtler et al. 2019; Szulzewsky et al. 2020).

Several questions remain about the biology of meningiomas: for example, does a subset of *NF2*-wild type meningiomas harbor alternative *NF2*-mutant-like mutations that ultimately activate similar pathways and oncogenic drivers, and do these tumors harbor and rely on oncogenic YAP activity? Even though the rarity of certain cancer subtypes often renders them infeasible for specific larger clinical trials, sometimes rare drivers of a complex disease can be informative of the overall biology of the disease.

In this study, we show that human *NF2*-wild type YAP1 fusion-positive meningiomas mimic *NF2*-mutant meningiomas in their global and YAP1-related gene expression signatures, suggesting that both tumor types harbor activated YAP signaling. We then utilize our RCAS/tv-a system for somatic cell gene transfer and show that intracranial expression of *YAP1-MAML2* results in a high penetrance of tumors in mice that resemble human meningiomas by histology and gene expression. We then show that YAP1-MAML2 exerts TEAD-dependent YAP activity that is resistant to inhibitory Hippo pathway signaling and can be targeted *in vitro* by pharmacological disruption of the YAP1-TEAD interaction. This elevated and unregulated YAP activity is sufficient to drive the formation of these tumors as we show that expression of constitutively active NLS-2SA-YAP1 on its own is also sufficient to induce very similar meningioma-like tumors in mice. In summary, our results suggest 1) that *YAP1-MAML2* is a causal oncogenic driver in pediatric NF2-wild type meningioma, 2) that *YAP1-MAML2* represents an alternative route of achieving de-regulated and oncogenic YAP activation in meningioma in addition to *NF2* loss, and 3) that de-regulated TEAD-dependent YAP activity is an oncogenic driver in *YAP1-MAML2*-fusion meningioma as well as NF2-mutant meningioma in general.

## Results

### Human NF2-wild type YAP1 fusion-positive meningiomas resemble NF2-mutant meningiomas by gene expression

We have previously shown that several YAP1 gene fusions found in other cancers exert de-regulated YAP1 activity that is insensitive to inhibitory Hippo pathway signaling (Szulzewsky et al. 2020). Considering that NF2, a potent tumor suppressor and upstream regulator of YAP1, is functionally lost or inactivated in a large percentage of meningiomas, we speculated that 1) activation of YAP1 signaling is present in *NF2*-mutant meningiomas and that 2) *NF2*-wild type YAP1 fusion-positive meningiomas also harbor YAP1 signaling and resemble *NF2*-mutant meningiomas on a gene expression level.

To determine how similar the gene expression patterns of human YAP1 fusion-positive (human YAP1fus) meningiomas are to the more common meningiomas, and specifically *NF2*-mutant (NF2mut) meningiomas, we analyzed the RNA-Seq data of 221 human meningioma samples from three different datasets (human YAP1fus (6 samples), NF2mut (Chr22 loss and/or presence of mutations in *NF2*; 104 samples), NF2wt_TKAS (presence of mutations in *TRAF7, KLF4, AKT1*, or *SMO1*, absence of Chr22 loss or mutations in *NF2;* 43 samples), NF2wt_NOS (absence of Chr22 loss or mutations in *NF2, TRAF7, KLF4, AKT1*, or *SMO1;* 68 samples)) and Pilocytic Astrocytoma (PA) samples (10 samples) (Patel et al. 2019; Prager et al. 2020; Sievers et al. 2020a; Maas et al. 2021), (Supplemental Table S1). Additional methylation classifier data was available for 42 tumors.

In order to create a 2D reference landscape, we used the Uniform Manifold Approximation and Projection (UMAP) dimensional reduction algorithm to cluster tumor samples based on their global gene expression patterns and found that the PA samples clustered distinctly from meningioma samples (Fig. 1A-B). We calculated the UMAP with the complete dataset and are only showing human samples. Additional UMAPs calculated with only the human samples show similar results and can be found in Supplemental Fig. S1A-B. We observed that NF2mut and NF2wt_TKAS tumors separated into distinct regions of the overall meningioma cluster (Fig. 1C-D). Within the NF2mut region, WHO grade 1 tumors mostly separated from WHO grade 3 tumors, whereas grade 2 tumors were distributed across the entire NF2mut population (Fig. 1B). For the tumors for which methylation classifier data was available we found that WHO grade 2 tumors that clustered with WHO grade 1 tumors were predominantly of the benign methylation subtype whereas grade 2 tumors that clustered with WHO grade 3 tumors were of the malignant subtype (Supplemental Fig. S1A). Human YAP1fus tumors clustered with WHO grade 1-2 NF2mut tumors of the benign methylation subtype (Fig. 1C-D, Supplemental Fig. S1A), indicating that they share a similar gene expression profile compared to these tumors.

**Figure 1:**
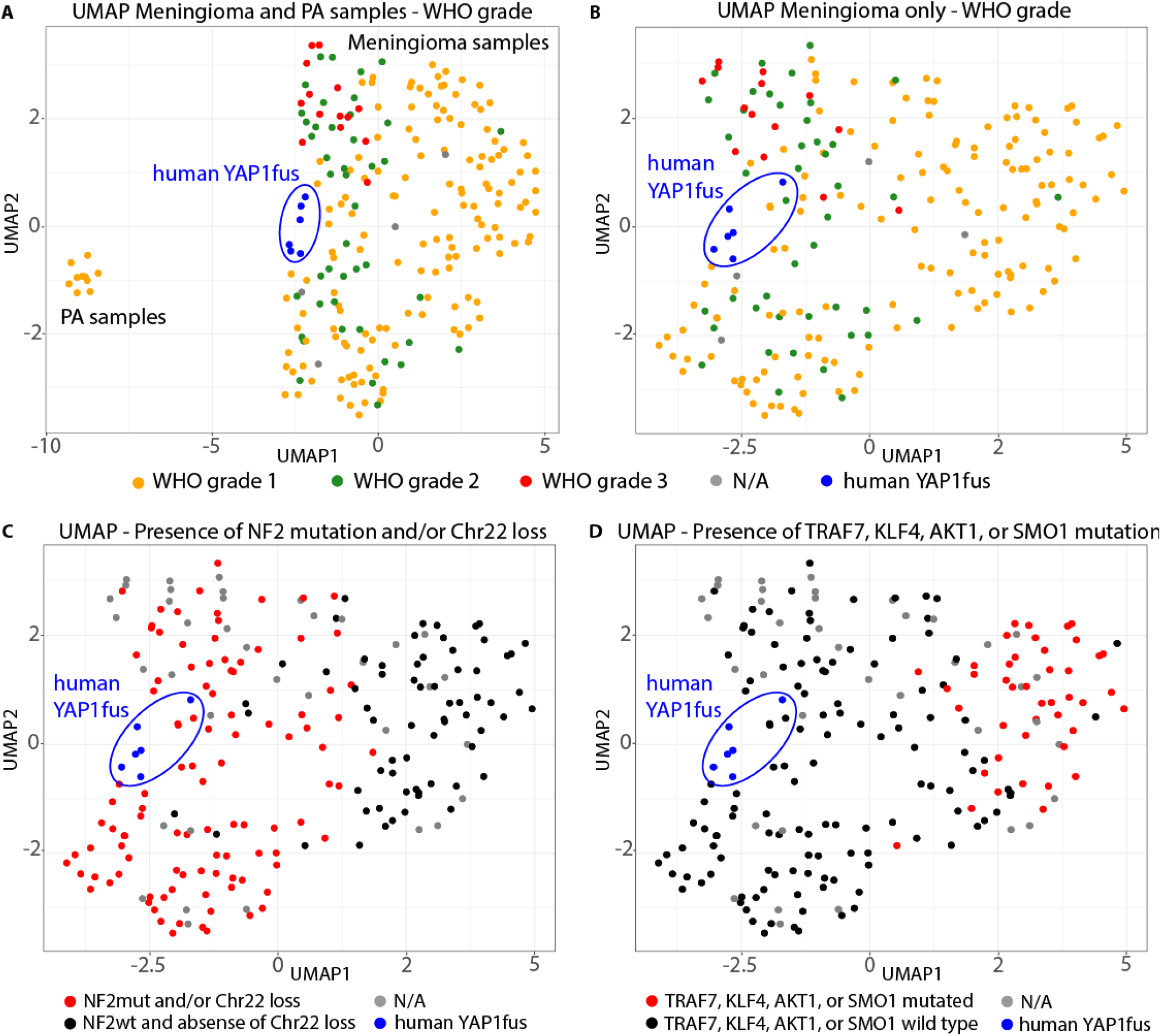
Human *NF2*-wild type YAP1 fusion-positive meningiomas resemble *NF2*-mutant meningiomas by gene expression. A-B)UMAP with RNA-Seq data of human meningioma samples (including YAP1 fusion-positive meningiomas) with (A) or without (B) Pilocytic Astrocytoma (PA) samples. Samples are colored in by WHO grade. C-D) Same UMAP as (B), with samples being colored in by the presence of (C) *NF2* mutations and/or Chr22 loss or (D) mutations in *TRAF7, KLF4, AKT1*, or *SMO1*.

We also performed hierarchical clustering of this data which showed that PAs clustered away from meningioma samples and that NF2mut and NF2wt_TKAS samples predominantly segregated apart from each other (Supplemental Fig. S1C). The hierarchical clustering, similar to the UMAP analysis above, places the human YAP1fus tumors with NF2mut tumors, again highlighting the similarity in the overall gene expression of these sample groups (Supplemental Fig. S1C).

Taken together, human YAP1 fusion-positive tumors show a similar gene expression pattern as more benign WHO grade 1-2 NF2mut meningiomas.

### Human YAP1 fusion-positive meningiomas and NF2-mutant meningiomas exert increased levels of YAP signaling

NF2 is an upstream regulator in the Hippo Signaling Pathway which inhibits the activity of YAP1. We therefore analyzed the expression of YAP1 downstream target genes in the different meningioma samples groups with known mutational status for *NF2, TRAF7, KLF4, AKT1*, or *SMO1* (human YAP1fus, NF2mut, NF2wt_TKAS).

We analyzed the expression of *NF2* and *YAP1*, as well as several direct YAP1 target genes (*CTGF, CYR61, ANKRD1, AMOTL2, CPA4, AJUBA, ANXA1, ANXA3*, and *CITED2*) in the different sample groups (Fig. 2A, Supplemental Fig. S2A). *NF2* expression was significantly lower in NF2mut tumors compared to human YAP1fus tumors. We observed high *YAP1* expression in all meningioma samples, whereas PA samples displayed a significantly lower expression. In line with these results, we observed that the YAP1 downstream target genes *CTGF, ANKRD1, AMOTL2*, and *CP4* were expressed at significantly higher levels in human YAP1fus tumors compared to NF2wt_TKAS and PA tumors but not to NF2mut tumors, suggesting that NF2mut and human YAP1fus tumors regulate these YAP1 targets at similar levels. We observed a similar non-significant trend for *CYR61*. By contrast, the expression of *AJUBA, ANXA1, ANXA3*, and *CITED2* was high in all meningioma subtypes but low in PA samples. These results suggest that both human YAP1fus and NF2mut tumors show higher expression levels of several canonical YAP1 target genes compared to both PA samples (that express only low levels of *YAP1*) and NF2wt_TKAS tumors (that express high levels of *NF2*).

**Figure 2:**
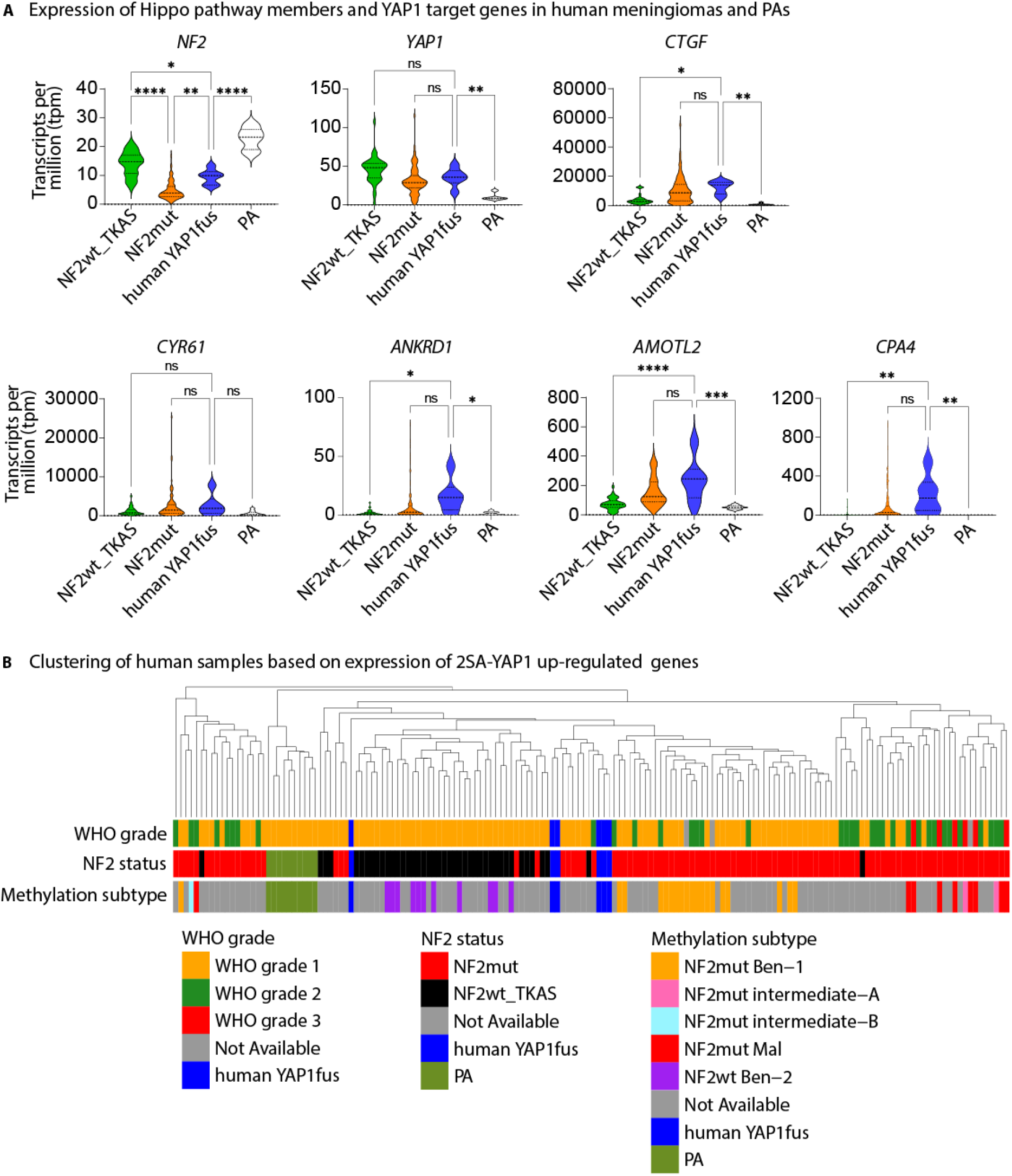
Human YAP1 fusion-positive meningiomas and *NF2*-mutant meningiomas exert increased levels of YAP signaling. A) Expression of *NF2, YAP1*, and selected YAP1 target genes (*CTGF, CYR61, ANKRD1, AMOTL2*, and *CPA4*) in human meningiomas (NF2wt_TKAS, NF2mut, humanYAP1fus) and PAs. B) Clustering of human samples based on the expression of 2SA-YAP1 up-regulated genes. Error bars show SEM. Analysis was done using ordinary one-way ANOVA (A). (*) P < 0.05; (**) P < 0.01; (***) P < 0.001; (****) P < 0.0001.

To analyze the expression of YAP1-regulated genes in the different Meningioma samples as well as in the PA samples on a broader scale, we generated a dataset of YAP1-regulated genes based on RNA-Seq data that we had previously generated from human neural stem cells expressing an activated version of YAP1 (S127/397A-(2SA)-YAP1; 1116 up- and 1501 down-regulated genes compared to GFP-expressing control cells) (Fig. 2B, Supplemental Fig. S2B-C, Supplemental Table S2) (Szulzewsky et al. 2020). We performed hierarchical clustering based on the expression of these 2SA-YAP1-regulated genes and found that PA samples clustered away from meningioma samples, indicating that meningiomas exhibit altered YAP signaling compared to PAs. NF2mut and NF2wt_TKAS tumors clustered mostly separate from each other. Human YAP1fus tumors clustered more closely with NF2mut tumors. Lower-grade (WHO grade 1-2 with benign-1 methylation subtype) and higher-grade (WHO grade 2-3 with intermediate or malignant methylation subtype) NF2mut meningiomas clustered separate from each other. We found that human YAP1fus tumors predominantly clustered more closely with lower-grade NF2mut tumors when clustering was based on the 1116 2SA-YAP1 up-regulated genes, but clustered with higher-grade NF2mut tumors based on 1501 2SA-YAP1 down-regulated genes.

Taken together, our results indicate that human YAP1fus meningiomas exert a YAP1-related gene expression signature that closely resembles that of NF2mut meningiomas.

### Forced expression of YAP1-MAML2 induces the formation of meningioma-like tumors in mice

To determine if the expression of YAP1-MAML2 (YM) is sufficient to cause the formation of meningioma-like tumors in mice, we cloned HA-tagged versions of YM into the RCAS retroviral vector. Two different structural variants of YM have been identified in pediatric *NF2*-wild type meningioma: the shorter variant retains only the first exon (amino acids 1-107) of the YAP1 sequence, whereas the longer variant retains exons 1-5 (amino acids 1-328). Both variants are fused to exons 2-5 (amino acids 172-1152) of MAML2. Since both variants of YM exceed the maximum capacity of the RCAS vector (around 2.5 kilobases), we generated two different truncated versions of the shorter YM variant for our *in vivo* studies (Supplemental Fig S3A-C). Truncated version 1 (YMv1) lacked part of the C-terminal sequence of MAML2 (lacking amino acids 885 – 1141 of wtMAML2), whereas truncated version 2 (YMv2) retained the C-terminus of MAML2 (lacking amino acids 321 – 569 of wtMAML2).

We utilized the RCAS/tv-a system for somatic cell gene transfer in combination with Nestin/tv-a (N/tv-a), to intracranially express the different YM constructs in Nestin-positive cells of p0-p3 neonatal pups. We have previously used the same system to study the oncogenic functions of other YAP1 gene fusions (Szulzewsky et al. 2020) and other oncogenic drivers (Ozawa et al. 2018). Immunohistochemical staining of cranial tissue from p0 neonatal N/tv-a pups showed strong Nestin staining in the meninges, confirming that these mice will support infection by RCAS vectors in the meninges and near the ventricles (Supplemental Fig. S3D). Intracranial expression of YMv2 in N/tv-a *Cdkn2a* wild type mice resulted in small lesions near the ventricles 150 days post-injection (Supplemental Fig. S3E-F). For modeling purposes, and to generate tumors in the life span of a mouse we used a *Cdkn2a*-deficient genetic background. *CDKN2A/B* loss is seen in a subset of NF2-mutant meningiomas but so far not in the nine YAP1 fusion tumors on record (Sievers et al. 2020a; Sievers et al. 2020b).

In humans, YAP1 fusion-positive meningiomas arise in different locations, both as extra-axial and intra-ventricular tumors (Ostrom et al. 2020). We determined if the intracranial expression of the two different truncated YM variants could induce tumor formation in different locations in N/tv-a *Cdkn2a* null neonatal mice (Fig. 3A-B; Supplemental Fig. S3G-H). We first wanted to determine if the two constructs are able to induce tumor formation. We performed injections into the brain parenchyma close to the ventricles and observed that both truncated variants of YM were able to induce meningioma-like tumors. Injection of YMv1 resulted in tumor formation in five out of twelve mice (41.7% penetrance), including one extra-axial, two intra-ventricular, and two extra-cranial tumors. In addition, three out of seven mice injected with YMv2 developed intracranial tumors (42.9% penetrance; one extra-axial tumor and two intra-ventricular tumors), that resembled tumors generated by the expression of YMv1. Because both truncated versions of YM were able to induce tumor formation, we then performed a second set of injections in which we injected YMv2 more superficially into the subarachnoid space of neonatal N/tv-a *Cdkn2a* null mice, since the likely cell-of-origin of extra-axial tumors is located within the meninges. We observed the formation of tumors in thirteen out of nineteen mice (68.4% penetrance), including six extra-axial and five intra-ventricular (centered in the lateral ventricle) tumors (Fig. 3A-B). Extra-axial tumors frequently eroded through the skull and also grew extra-cranially. Several mice exhibit the formation of co-occurring tumors in different locations. We monitored the growth of two extra-axial tumors by MRI and also performed weighed T1 MRIs with and without administration of contrast reagent. The tumors were diffusely contrast enhancing, similar to what is observed with human meningiomas (Fig. 3C, Supplemental Fig. S3I).

**Figure 3:**
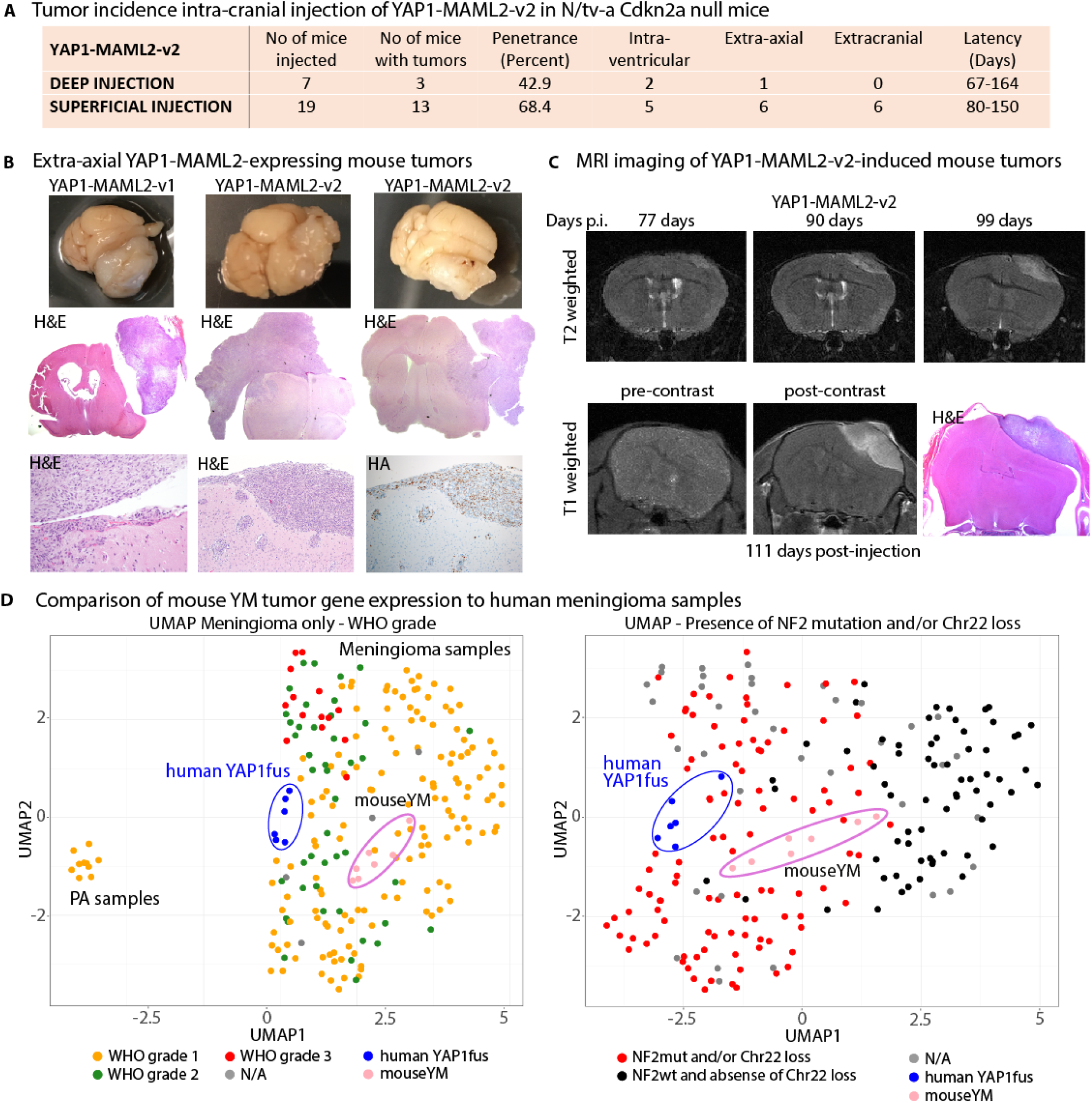
Forced expression of YAP1-MAML2 induces the formation of meningioma-like tumors in mice. A) Table showing the tumor incidence upon injection of YAP1-MAML2-(YM)-v2 in N/tv-a Cdkn2a null mice. B) Upper panel: gross-morphological pictures of one YM-v1-induced and two YM-v2-induced mouse meningiomas. Middle panel: corresponding H&E images of the tumors in the upper panel. Lower panel: higher magnification H&E stainings of YM-v2-induced tumors showing meningothelial features (left) and an example of meningioangiomatosis (middle). Tumors showed strong HA-positivity (right). C) Upper panel: T2w MRI images taken 77, 90, or 99 days post-injection. Lower panel: T1w MRI images pre- (left) or post-contrast (middle) taken 111 days post-injection. H&E staining of the same tumor (right). D) UMAP of mouseYM, human meningioma and PA samples. Samples are colored in by WHO grade (left) or by the presence of *NF2* mutations and/or Chr22 loss (right).

The extra-axial tumors were generally well-circumscribed spindle cell tumors resembling meningiomas on histomorphology and were located in the meninges (Supplemental Fig. S3J). A subset of these tumors demonstrated a meningioangiomatosis type growth pattern with a downward spread along perivascular spaces (Fig. 3B). Histologically, the tumors were variably biphasic, with a predominantly compact spindle cell component and a less common loosely arranged spindle cell component with a myxoid-like background (Supplemental Fig. S3J-N). The compact spindle cell component was generally arranged in fascicles, imparting a fibrous-type appearance. Less commonly seen were lobules, reminiscent of traditional meningothelial meningioma appearance (Fig. 3B). Cytologically, the tumor cells were somewhat enlarged and had elongated nuclei with some nuclear irregularity and hyperchromasia (Supplemental Fig. S3O). Occasionally the cells were more pleomorphic (Supplemental Fig. S3P). There was variable mitotic activity found in these tumors, ranging up to 80 mitoses per mm^2^ for YMv2-derived tumors, corresponding to atypical meningioma WHO grade 2, and up to 403 mitoses per mm^2^ for YMv1-derived tumors, corresponding to an anaplastic meningioma WHO grade 3 (Supplemental Fig. S3Q-R). All tumors showed positive immunohistochemical staining for the HA tag (Fig. 3B). Several tumors displayed focal positive staining for EMA, and YMv2 tumors displayed similar *EMA/Muc1* gene expression compared to human meningiomas (Supplemental Fig. S3R-S). The tumors/tumor cells stained positive for Vimentin, but were negative for Synaptophysin, GFAP, and OLIG2 (Supplemental Fig. S3U). We observed areas that showed positive staining for OLIG2, however co-immunofluorescence stainings showed that there was no overlap between the tumor cells that stained positive for the HA tag and OLIG2-positive cells, suggesting that the OLIG2-positive cells are entrapped non-transformed glia cells (Supplemental Fig. S3V-W).

We did not observe any tumor formation upon intracranial injection of RCAS vectors encoding either wtYAP1 or wtMAML2 (Supplemental Fig. S3H). Due to the size limitations of the RCAS vector we expressed truncated versions of wtMAML2 that shared the same truncations as YMv1 and YMv2 (Supplemental Fig. S3A-B).

These results suggest that the expression of YAP1-MAML2 is sufficient to cause tumor formation from Nestin-expressing cells in the subarachnoid and ventricular spaces and that YAP1 fusions are the likely oncogenic drivers in human YAP1 fusion-positive meningiomas.

### Murine YAP1-MAML2-driven tumors resemble human YAP1fus and NF2mut meningiomas by gene expression

To determine if the YAP1-MAML2-expressing mouse tumors resemble human YAP1fus and NF2mut meningiomas by gene expression, we performed RNA-Seq on seven of our YMv2-driven mouse tumors (mouseYM; three extra-axial tumors, three extra-cranial tumors, one intra-ventricular tumor). To compare the mouse and human samples, we converted the mouse gene symbols to human gene symbols and only kept genes that are present in both species, leaving us with 16895 unique genes.

We then again used UMAP and found that the mouseYM samples clustered with NF2mut meningiomas, indicating that they express a similar gene expression profile (Fig. 3D, Supplemental Fig. S3X-Y). We again performed hierarchical clustering based on the expression of 2SA-YAP1-regulated genes (1116 up- and 1501 down-regulated genes) and found that mouseYM tumors clustered closely with human YAP1fus and NF2mut meningiomas (Supplemental Fig. S3Z-AA). Similarly, we observed that mouseYM tumors expressed several YAP1 target genes at similar levels compared to human YAP1fus and NF2mut meningiomas (Supplemental Fig. S3AB).

Taken together, our results show that mouseYM meningioma-like tumors resemble human YAP1fus and NF2mut meningiomas by gene expression, both, when considering the overall and the YAP1-related gene expression.

### YAP1-MAML2 is constitutively localized to the nucleus and insensitive to Hippo pathway-mediated inhibition

We have previously shown that several YAP1 fusion proteins are constitutively localized to the nucleus - mediated by an NLS in the sequences of the C-terminal fusion partners – and that this nuclear localization is essential to the oncogenic abilities of the YAP1 fusion proteins (Szulzewsky et al. 2020).

We performed immunofluorescence (IF) stainings of HEK293 (cultured at high-density conditions) expressing HA-tagged versions of wtYAP1, Y(e1)M, or wtMAML2 (Fig. 4A). As previously reported, wtYAP1 staining localized mostly to the nucleus at low cell densities but was excluded from the nucleus at high cell densities, whereas YM and wtMAML2 displayed constitutive nuclear staining at all cell densities. In addition, HA-tag IHC stainings of YMv1 and YMv2 mouse tumors *in vivo* revealed strong nuclear localization of the fusion protein (Fig. 4B, Supplemental Fig. S4A). These results suggest that the fusion of the YAP1 sequence to MAML2 prevents it from being excluded from the nucleus upon high cell densities.

**Figure 4:**
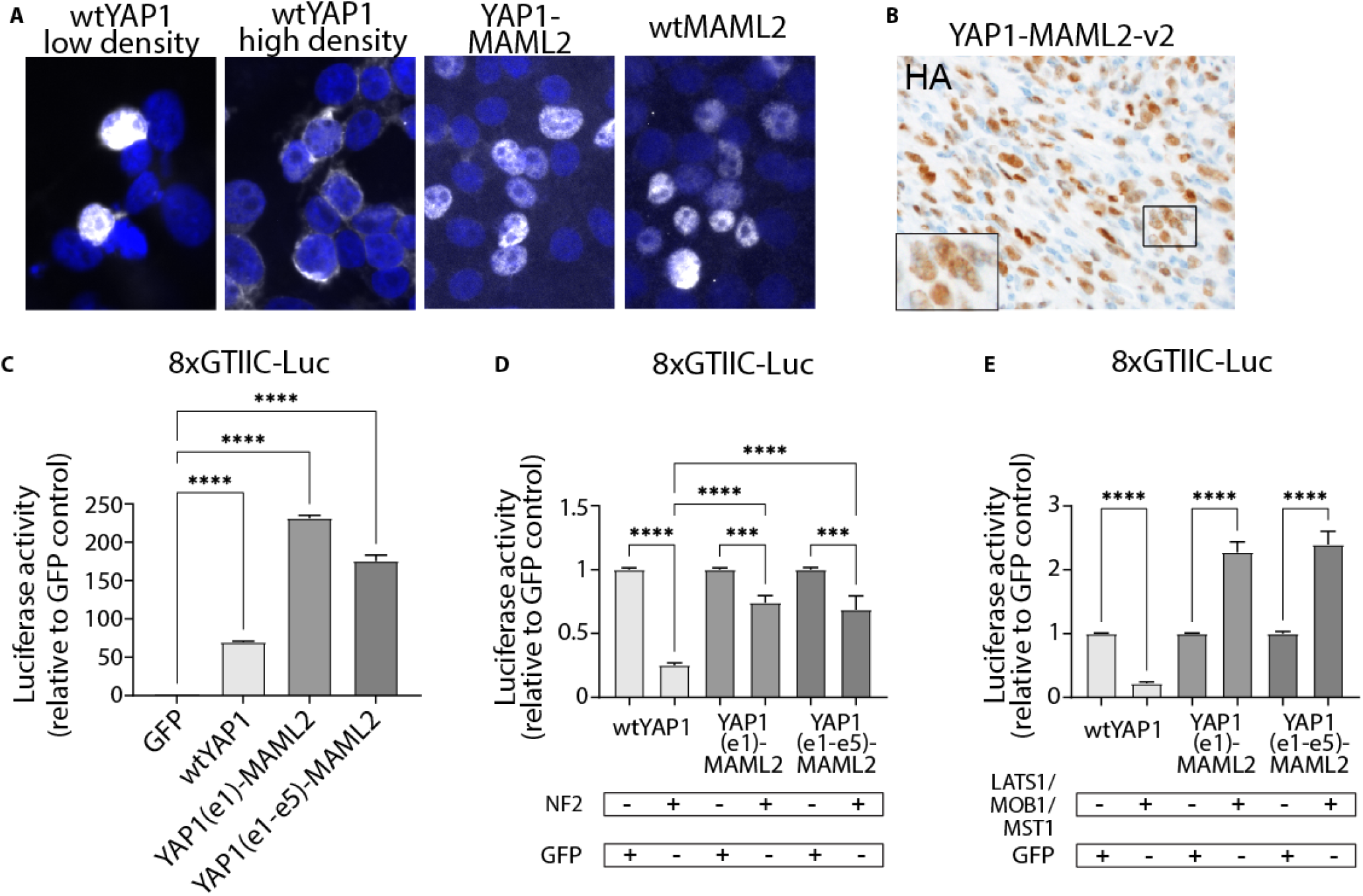
YAP1-MAML2 is constitutively localized to the nucleus and is insensitive to Hippo pathway-mediated inhibition. A) HA IF stainings (Hoechst counter stain) of confluent (except wtYAP1 low density) HEK293 cells expressing wtYAP1, YAP1(e1)-MAML2, or wtMAML2. B) HA IHC staining of a YMv2 mouse tumor demonstrates nuclear localization of YM *in vivo*. C) Activity of GFP, wtYAP1, Y(e1)M, and Y(e1-e5)M in the YAP1-responsive GTIIC-Luc reporter assay (n = 3). D) Effect of additional *NF2* coexpression on the YAP activity of wtYAP1 (n = 8), Y(e1)M (n = 8) or Y(e1-e5)M (n = 6) in the GTIIC-Luc reporter assay. E) Effect of additional LATS1/MOB1/MST1 co-expression on the YAP activity of wtYAP1 (n = 10), Y(e1)M (n = 10) or Y(e1-e5)M (n = 4) in the GTIIC-Luc reporter assay. Error bars show SEM. Analysis was done using ordinary one-way ANOVA (C,D,E). (***) P < 0.001; (****) P < 0.0001.

Similar to the YAP1 fusions that we previously analyzed, YM retains the TEAD binding domain near the N-terminus of wild type YAP1, suggesting that YAP1-MAML2 is able to exert YAP activity. To analyze the baseline YAP activity of the different proteins, we transiently transfected HEK293 cells at sub-confluency cell densities with the YAP1-responsive 8xGTIIC-Luc YAP1 reporter plasmid and either GFP (control), wtYAP1, or YM. In addition to the shorter YM variant (Y(e1)M, retaining only exon1 of YAP1), we also analyzed the longer YM variant (Y(e1-e5)M, retaining exons1-5 of YAP1) (Supplemental Fig. S3A). We observed that wtYAP1 as well as both YM variants significantly activated the YAP1-responsive reporter compared to GFP control cells (p < 0.0001 for all) (Fig. 4C). Variable amounts of the steady state protein between constructs were observed that do not correlate with YAP activity (Supplemental Fig. S4B). These results show that YM exerts YAP transcriptional activity.

We determined the effect of additional co-expression of *NF2* or the Hippo pathway proteins LATS1, MST1, and MOB1 (compared to GFP control) using GTIIC-Luc reporter assays in transiently transfected HEK293 cells. The activity of wtYAP1 and YM was significantly reduced (padj < 0.0001) by co-expression *NF2*, however the YAP activity of YM was significantly less affected (padj < 0.0001) (Fig. 4D). In turn, while the activity of wtYAP1 was significantly reduced (padj < 0.0001) upon co-expression of LATS1/MST1/MOB1, the YAP activity of YM was significantly increased (padj < 0.0001) (Fig. 4E). These results suggest that Hippo signaling actually promotes the activity of YM, rather than inhibiting it, even though the underlying mechanism remain currently unknown. Of note, both the short and the long variant of YM behaved similarly, even though the longer Y(e1-e5)M variant retains several of the serine residues targeted by LATS1/2 (including S127 that is lost in Y(e1)M). This indicates that the strong nuclear localization of the MAML2 protein outweighs the Hippo pathway-mediated cytoplasmic retention of YAP1. We had previously observed similar results with other YAP1 fusions (such as YAP1-MAMLD1 and YAP1-FAM118B) that also retain several of the serine residues targeted by LATS1/2 (Szulzewsky et al. 2020).

Taken together, our results show that – in contrast to wtYAP1 – the YAP activity of YAP1-MAML2 is not inhibited by Hippo pathway signaling.

### The transcriptional program induced by YAP1-MAML2 is dependent on the interaction with TEAD transcription factors

We have previously shown that several YAP1 fusions rely on the interaction with TEAD transcription factors to exert their pro-oncogenic transcriptional programs (Szulzewsky et al. 2020). To analyze the transcriptional programs induced by YM on a larger scale and determine to what extend it relies on the interaction with TEADs, we transfected HEK cells with RCAS plasmids containing either wtYAP1, Y(e1)M, S94A-Y(e1)M, or GFP, isolated RNA 48hrs post-transfection, and performed RNA-Seq. Principal Component Analysis (PCA) clearly separated the different sample groups (Fig. 5A). YM-expressing cells exhibited the greatest number of DEGs compared to GFP-expressing cells and up-regulated the expression of the direct YAP1 target genes *CTGF, CYR61, ANKRD1*, and *AMOTL2*, more strongly than wtYAP1-expressing cells (Fig. 5B-C, Supplemental Fig. S5A-E).

**Figure 5:**
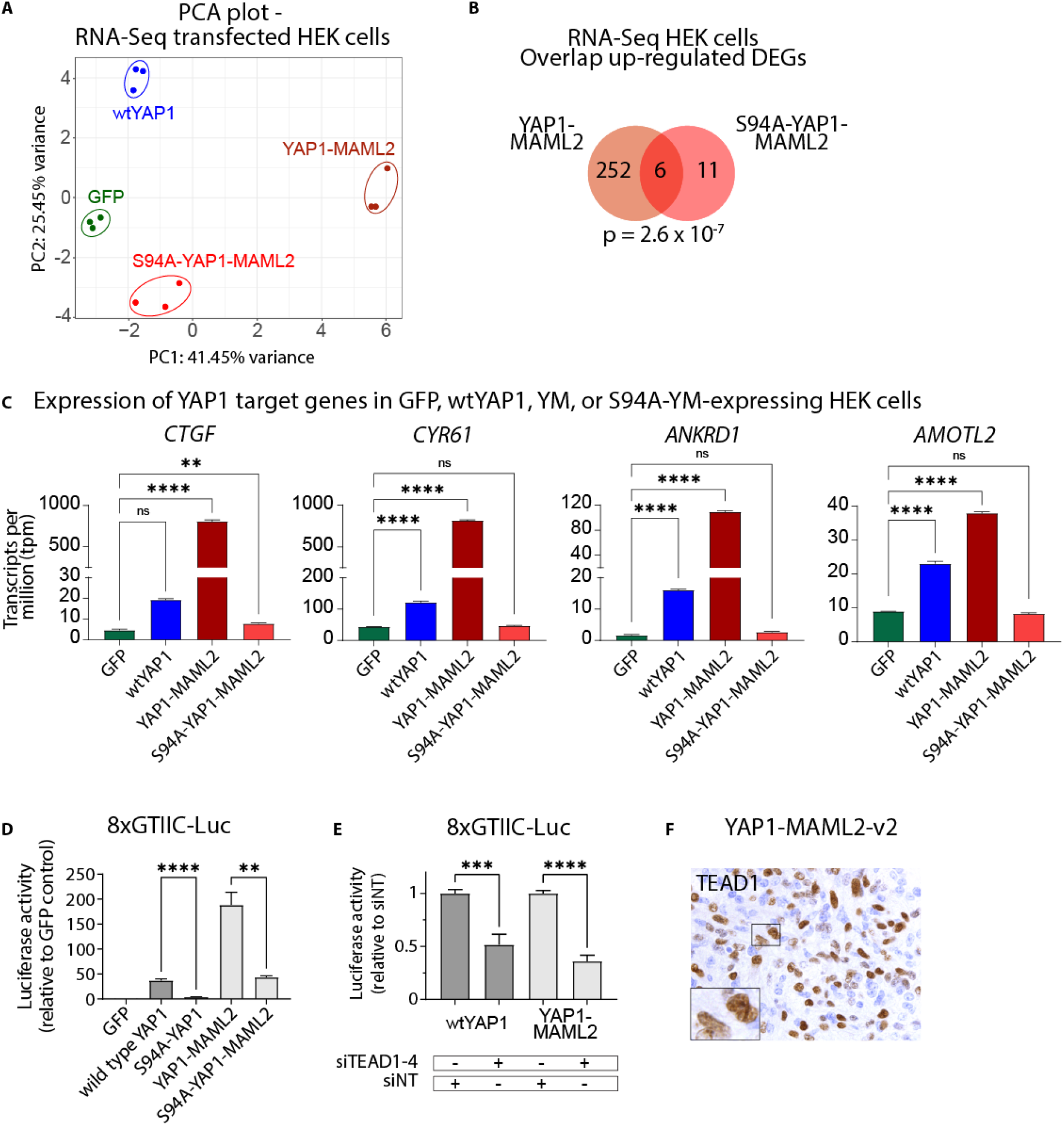
The transcriptional program induced by YAP1-MAML2 is dependent on the interaction with TEAD transcription factors. A) PCA plot of RNA-Seq samples. B) Venn diagram showing the overlap of up-regulated DEGs (compared to GFP-expressing cells) between YM- and S94A-YM-expressing cells. C) Expression of the YAP1 target genes *CTGF, CYR61, ANKRD1*, and *AMOTL2* in GFP-, wtYAP1-, YM-, or S94A-YM-expressing HEK cells. D) YAP activity of S94A mutant wtYAP1 and YM in the GTIIC-Luc reporter assay (n = 4 each). E) Combined knockdown of TEAD1–4 leads to reduced YAP activity of wtYAP1 and YM in the GTIIC-Luc reporter assay (n = 6). F) TEAD1 IHC staining of a YM-v2 mouse tumor. Error bars show SEM. Analysis was done using ordinary one-way ANOVA (C) or two-tailed t-test (D,E). (**) P < 0.01; (***) P < 0.001; (****) P < 0.0001.

By contrast, S94A-YM (a point mutant that is unable to bind to TEAD transcription factors) did not recapitulate the gene expression changes caused by YM and did not significantly induce the expression of specific YAP1 target genes (Fig. 5B-C, Supplemental Fig. S5D-E). Similarly, we observed that the S94A-mutant versions of wtYAP1 (p < 0.0001) and YM (p = 0.0013) displayed a significantly reduced ability to activate the 8xGTIIC-Luc reporter compared to their unmutated counterparts (Fig. 5D, Supplemental Fig. S4B), indicating that the interaction with TEAD transcription factors is crucial for their functions. The combined knockdown of TEADs 1-4 also resulted in a significantly reduced ability of wtYAP1 (p = 0.001) and YM (p < 0.0001) to activate the 8xGTIIC-Luc reporter (Fig. 5E).

Taken together, our results show that the transcriptional activity of YAP1-MAML2 significantly relies on the interaction with TEAD transcription factors.

### The interaction with TEAD transcription factors is necessary for the oncogenic activity of YAP1-MAML2 in vivo

We have previously shown that the interaction with TEAD transcription factors is essential for the oncogenic functions of other YAP1 gene fusions and that this interaction can be blocked by small molecule inhibitors, such as Verteporfin (Liu-Chittenden et al. 2012), which in turn leads to a reduction in the viability of tumor cells and *ex vivo*-cultured tumor slices (Szulzewsky et al. 2020).

We detected robust expression of all four TEADs in all meningioma subtypes (Supplemental Fig. S5F). In addition, we found that a large percentage of cells in our mouseYM tumors stained positive for TEAD1, whereas only a minority of cells showed TEAD1 staining in naïve brain sections (Fig. 5F, Supplemental Fig. S5G).

To determine whether the interaction with TEAD transcription factors is necessary for the oncogenic functions of YM, we intracranially expressed S94A-YMv2 in N/tv-a *Cdkn2a* null mice (Supplemental Fig. S5H). We found that compared to YMv2, S94A-YMv2 showed a significantly reduced oncogenic capacity (Fig. 6A). None of the 16 mice expressing RCAS-S94A-YMv2 developed tumors, indicating that the interaction with TEADs is necessary for the ability of YAP1-MAML2 to cause tumor formation.

**Figure 6:**
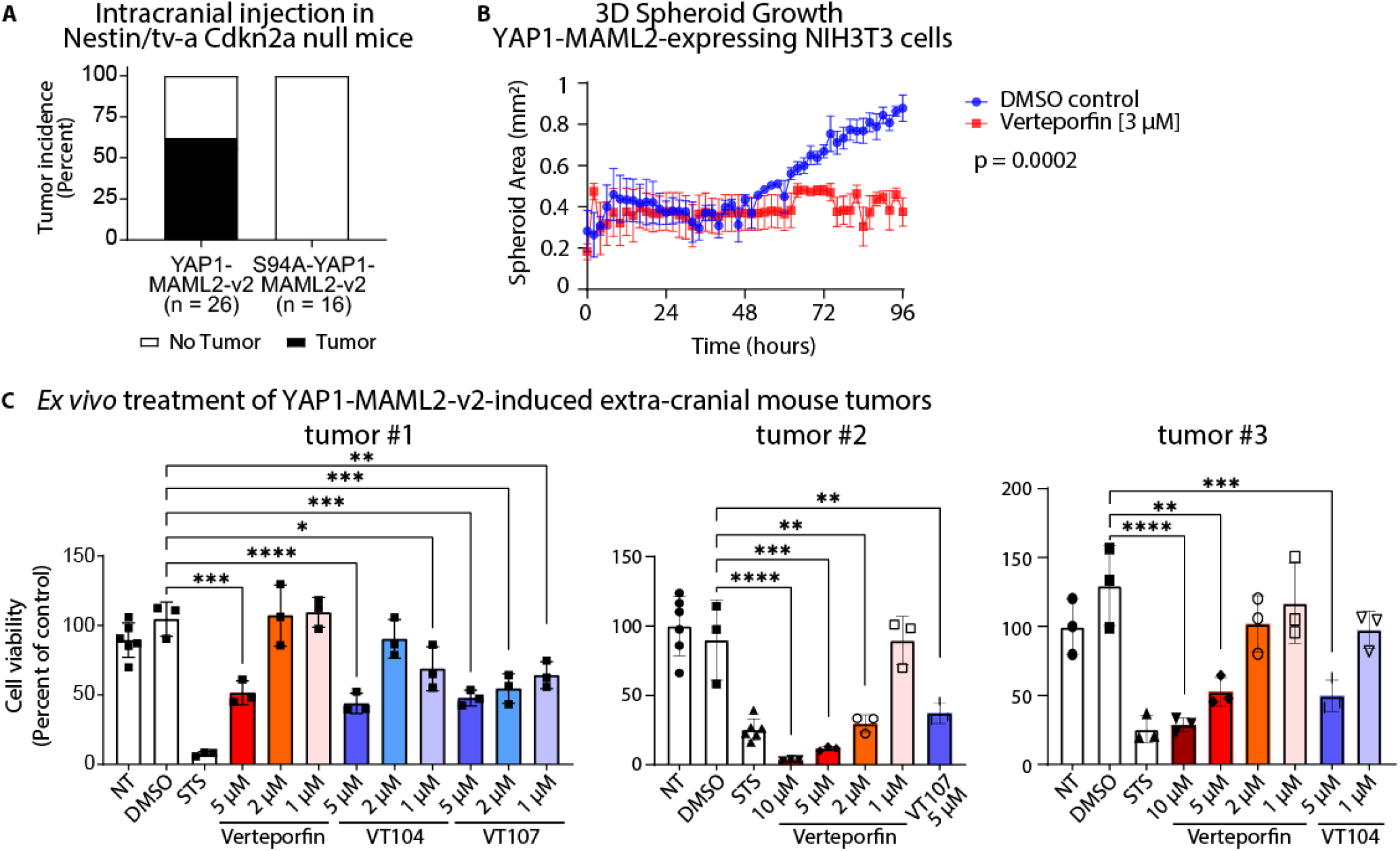
The interaction with TEAD transcription factors is necessary for the oncogenic activity of YAP1-MAML2 in vivo. A) Tumor incidence upon injection of S94A-YM-v2 into N/tv-a Cdkn2a null mice. B) Spheroid growth of YM-expressing NIH3T3 cells when treated with VP or DMSO only (n = 3). C) Viability of YM-v2 organotypic mouse tumor cuboids (taken from three separate extra-cranial tumors) after no treatment (NT) or treatment with either DMSO, Staurosporine (STS), Verteporfin, or VT104/VT107. Error bars show SEM (B) or SD (C). Analysis was done using ordinary two-way ANOVA (B) or ordinary one-way ANOVA (C). (*) P < 0.05; (**) P < 0.01; (***) P < 0.001; (****) P < 0.0001.

Several pharmacological inhibitors that can disrupt the interaction between YAP1 and TEADs are currently being evaluated in the pre-clinical setting (Pobbati and Hong 2020). To test if the oncogenic YAP activity of YM can be pharmacologically inhibited, we lentivirally transduced NIH3T3 cells to stably express either wtYAP1, wtMAML2, or YM. In 3D culturing conditions neither untransduced control cells, wtYAP1-expressing, nor wtMAML2-expressing cells were able to grow into spheroids, likely due to high contact inhibition (Holley and Kiernan 1968), whereas YM-expressing cells were able to grow into spheroids (Fig. 6B Supplemental Fig. S6A-C). Treatment with 3 μM Verteporfin inhibited the spheroid growth of YM-expressing cells (p < 0.0001) (Fig. 6B), accompanied by a significant down-regulation of the YAP1 downstream targets *Ctgf* and *Cyr61* (Supplemental Fig. S6D), while Verteporfin treatment did not affect the growth of either untransfected, wtYAP1- or wtMAML2-expressing cells (Supplemental Fig. S6A-C).

Lastly, we established tumor cuboids from four individual extracranial YMv2 tumors (generated in N/tv-a *Cdkn2a* null mice) and treated them with different concentrations of Verteporfin, as well as two additional TEAD inhibitors (Tang et al. 2021), to test if pharmacological disruption of the YAP1-TEAD interaction can inhibit the viability of YM-driven tumors. We observed a significant and dosedependent reduction of the viability of YMv2 tumor cuboids treated with either Verteporfin, VT104, or VT107, compared to untreated or DMSO-treated tumor cuboids (Fig. 6C, Supplemental Fig. S6E-F).

These results suggest that the interaction with TEAD transcription factors is necessary for the oncogenic functions of YAP1-MAML2 and that pharmacological disruption of this interaction is – at least *in vitro* and *ex vivo* – sufficient to inhibit the growth and viability of YAP1-MAML2-driven tumor cells.

### Expression of constitutively activated YAP1 itself is sufficient to cause the formation of meningioma-like tumors in mice

The above data suggests that the YAP activity generated by the YM fusion is necessary for tumor formation but does not address whether YAP activity alone is sufficient to induce meningioma-like tumors in mice.

We and others have previously shown that the introduction of two separate point mutations into the sequence of YAP1 (S127/397A-(2SA)-YAP1) is sufficient to de-regulate its activity (Zhao et al. 2007; Zhao et al. 2010; Szulzewsky et al. 2020). The high frequency of inactivating *NF2* mutations in human meningiomas suggests a functional linkage to de-regulated YAP activity in these tumors, however, activating point mutations in YAP1 have not been identified in meningiomas so far. To assess if we could use activated YAP1 as a surrogate for *NF2* loss and to induce similar tumors, we analyzed RNA-Seq data to compare the effects of *NF2* loss (sgNF2, CRISPR-Cas9-mediated knockout of *NF2*) and expression of 2SA-YAP1 in U5 human neural stem cells (compared to untreated control cells) (Szulzewsky et al. 2020; O’Connor et al. 2021). Both conditions shared a highly significant overlap in their DEGs (396 overlapping up-regulated and 445 down-regulated DEGs, p < 10^-314^ for both comparisons) and similarly regulated several direct YAP1 downstream target genes (Supplemental Fig. S7A-D). This data shows that *NF2* loss and the expression of de-regulated 2SA-YAP1 induce similar transcriptomic changes, suggesting that we can use activated 2SA-YAP1 as a surrogate for *NF2* loss.

For our in vivo experiments we used a version of 2SA-YAP1 containing an additional N-terminal nuclear localization sequence (NLS) that we have previously shown to possess an increased nuclear localization (Szulzewsky et al. 2020) (Supplemental Fig. S3B). To test if elevated and de-regulated YAP activity alone is sufficient to cause the formation of meningioma-like tumors in mice, we injected RCAS viruses encoding an HA-tagged version of NLS-2SA-YAP1 superficially into the subarachnoid space of neonatal N/tv-a *Cdkn2a* null mice. We chose a *Cdkn2a* null background to ensure more rapid tumor growth and because *CDKN2A/B* loss is observed in a subset of human NF2mut meningiomas (Sievers et al. 2020b).

We observed the formation of tumors in twenty-nine out of thirty mice, including seventeen mice with extra-axial tumors (Fig. 7A-B, Supplemental Fig. S7E-I). Extra-axial tumors frequently invaded into bone and scalp soft tissue and eroded through the skull and grew also extra-cranially (Supplemental Fig. S7E-L). Most mice exhibited the formation of co-occurring tumors in different locations, such as additional tumors centered in the lateral ventricles (Supplemental Fig. S7M-N). The histopathologic features of NLS-2SA-YAP1 tumors were similar to that of YM-induced tumors, including the presence of a variable biphasic spindle cell appearance with occasional nuclear pleomorphism and enlarged nucleoli (Supplemental Fig. S7O-P). Brain invasion and/or a meningioangiomatosis type growth pattern was seen in a subset of tumors (Supplemental Fig. S7E, Q-R). Variable mitotic activity was also identified (Supplemental Fig. S7S). All tumors showed positive immunohistochemical staining for the HA tag and for TEAD1 (Supplemental Fig. S7T-U). We again monitored the growth of two extra-axial tumors by MRI. The tumors were diffusely contrast enhancing, similar to what we observed for the YM-induced tumors, and what is observed with human meningiomas (Fig. 7B).

**Figure 7:**
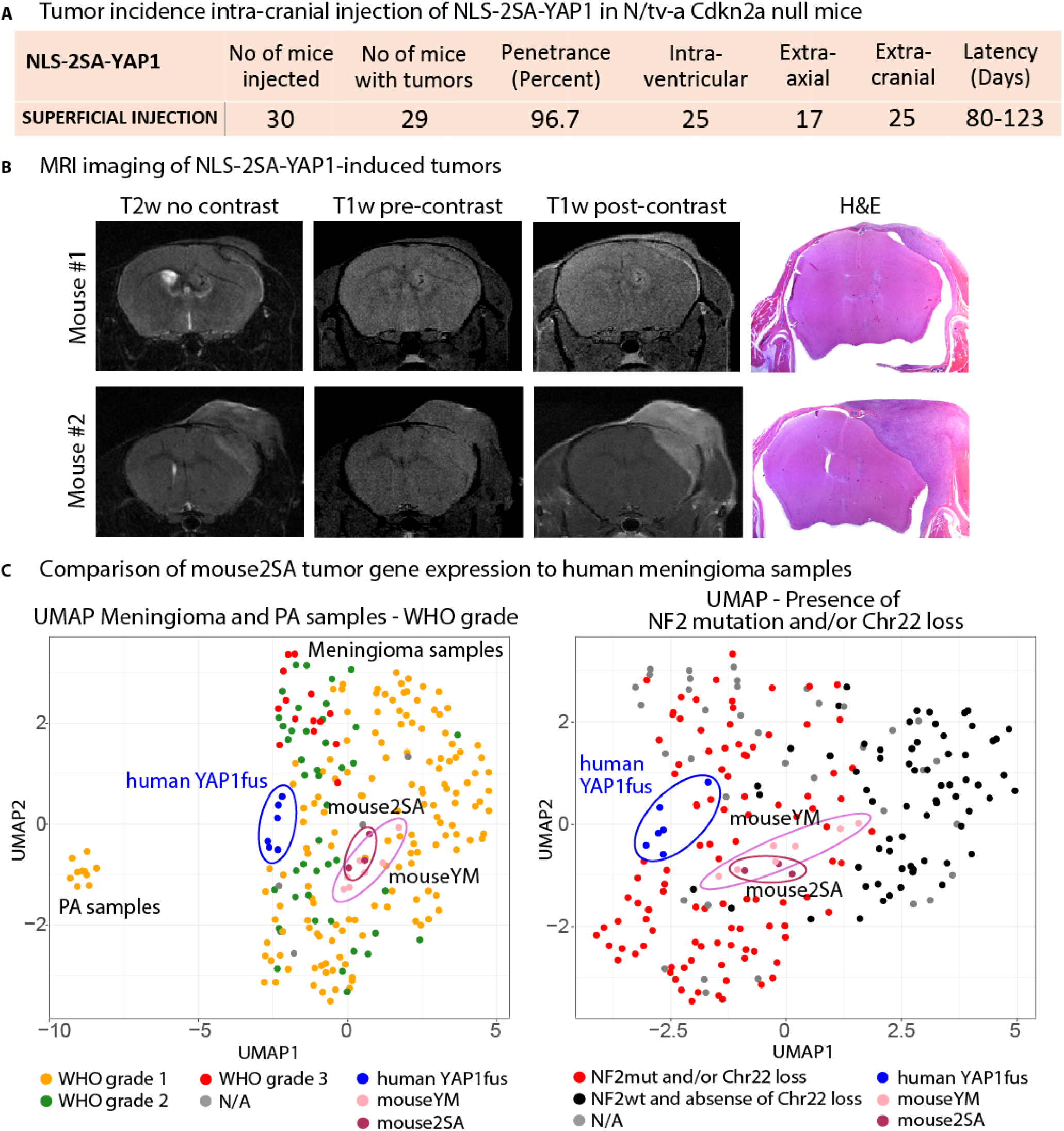
Expression of constitutively activated YAP1 itself is sufficient to cause the formation of meningioma-like tumors in mice. A) Table showing the tumor incidence upon injection of NLS-2SA-YAP1 in N/tv-a Cdkn2a null mice. T2w MRI images, T1w MRI images pre- or post-contrast, and H&E stainings of two NLS-2SA-YAP1-induced mouse tumors. D) UMAP of mouse2SA, mouseYM, human meningioma and PA samples. Samples are colored in by WHO grade (left) or by the presence of *NF2* mutations and/or Chr22 loss (right).

We extracted total RNA from three NLS-2SA-YAP1-driven mouse meningioma-like tumors (mouse2SA; one extra-axial tumor and two extra-cranial tumors) and performed RNA-Seq on these three samples to compare their gene expression to human meningiomas and mouseYM meningioma-like tumors. Based on the overall gene expression, mouse 2SA tumors clustered with human NF2mut meningiomas and mouseYM tumors, away from PAs (Fig. 7C, Supplemental Fig. S7V), indicating that these tumors resemble the overall gene expression pattern of human NF2mut meningiomas. In addition, hierarchical clustering based on the expression of 2SA-YAP1-regulated genes (1116 up- and 1501 down-regulated genes) showed that mouse2SA tumors clustered closely with human YAP1fus, mouseYM, and NF2mut meningiomas (Supplemental Fig. S7W-X). Furthermore, mouse2SA tumors showed increased expression of several direct YAP1 target genes, compared to PA samples (Supplemental Fig. S7Y). Finally, we again established tumor cuboids from five individual extra-cranial NLS-2SA-YAP1-driven mouse meningioma-like tumors and treated them ex vivo with different concentrations of either Verteporfin, VT104, or VT107. Similar to our results with YMv2-induced mouse tumors, we again observed a dose-dependent decrease in the viability of the tumor cuboids (Supplemental Fig. S7Z).

Taken together, our results show that the YAP activity generated by either the expression of the YAP1-MAML2 fusion or by constitutively active non-fusion YAP1 is sufficient to induce the formation of meningioma-like tumors in mice. Both tumor types resemble human meningiomas by histology and express a similar gene expression profile compared to human NF2mut meningiomas.

## Discussion

NF2/Merlin is a potent tumor suppressor, regulating the activity of the transcriptional co-activator and oncogene YAP1 via the Hippo signaling pathway (Petrilli and Fernandez-Valle 2016). Heterozygous deletion of chromosome 22 and additional functional inactivation of the remaining *NF2* gene copy occurs in around half of meningiomas (Riemenschneider et al. 2006), indicating that the de-regulation of YAP activity may play an important role in the pathobiology of these tumors. Mutations in *TRAF7, KLF4, AKT1*, and *SMO* have been identified in a subset of *NF2*-wild type tumors (Clark et al. 2013), however, the causal oncogenic drivers in tumors that do not harbor mutations in any of these genes remain largely unknown.

Recently, YAP1 fusions, first and foremost YAP1-MAML2, have been identified in a subset of pediatric *NF2*-wild type meningiomas (Sievers et al. 2020a). Even though these fusion cases are rare events, their occurrence in itself is informative about the underlying biology and the role of YAP signaling in meningioma. In this study, we show that 1) *YAP1-MAML2* is a causal oncogenic driver in pediatric NF2-wild type meningioma, 2) *YAP1-MAML2* represents an alternative route of achieving de-regulated and oncogenic YAP activation in meningioma in addition to *NF2* loss, and 3) de-regulated TEAD-dependent YAP activity is an oncogenic driver in *YAP1-MAML2*-fusion meningioma as well as NF2-mutant meningioma in general.

The expression of *YAP1-MAML2* is not only found in a subset of human meningiomas, but using the RCAS/tv-a system we can show that it is also sufficient to cause tumor formation in mice, suggesting that this fusion is the likely oncogenic driver in *YAP1-MAML2*-positive pediatric *NF2*-wild type meningiomas. This is in line with findings on other YAP1 gene fusions in several cancer types (Pajtler et al. 2019; Szulzewsky et al. 2020; Szulzewsky et al. 2021). The oncogenic effect of the fusion protein appears to be largely due to unregulatable YAP activity since we can show that 1) the YAP activity of the YAP1-MAML2 fusion protein is resistant to inhibitory Hippo signaling, 2) very similar tumors can be induced by constitutively-active non-fusion YAP1 (NLS-S127/397A-YAP1) constructs alone, and 3) genetic ablation of the YAP activity of YAP1-MAML2 (by S94A mutation) blocks its ability to form tumors.

Our hypothesis that YAP1-MAML2 represents an alternative route of achieving de-regulated and oncogenic YAP activation in addition to *NF2* loss is further supported by gene expression data showing that human YAP1 fusion-positive meningiomas harbor a gene expression signature that resembles *NF2*-mutant meningiomas, both on a global level as well as when specifically focusing on YAP1-regulated genes. Both *NF2*-mutant and YAP1 fusion-positive meningiomas express several YAP1 target genes (such as *CTGF, CYR61, AMOTL2, ANKRD1, CPA4*) at higher levels compared to *NF2*-wild type meningiomas and PAs, suggesting that both types of mutation similarly lead to an activation of YAP signaling. These findings are in line with previous studies that found increased YAP activity in *NF2*-mutant meningioma tumors and/or cell lines (Striedinger et al. 2008; Baia et al. 2012; Tanahashi et al. 2015).

YAP1 is a transcriptional co-activator that does not directly bind DNA, but functions through the interaction with other transcription factors, primarily TEADs (Zhao et al. 2008; Stein et al. 2015). The disruption of the interaction between YAP1 and TEADs (e.g., by introducing an S94A mutation into the YAP1 sequence) results in a severely reduced functionality of YAP1. Likewise, we show that the functionality of YAP1-MAML2 also largely relies on its interaction with TEADs, since an S94A-mutant of YAP1-MAML2 was unable to recapitulate the transcriptional changes induced by unmutated YAP1-MAML2 and was furthermore unable to cause tumor formation in vivo. These results are in line with previous findings on other YAP1 gene fusions (Pajtler et al. 2019; Szulzewsky et al. 2020; Szulzewsky et al. 2021). Moreover, we also show that pharmacological inhibition of the interaction between YAP1-MAML2 and TEADs by Verteporfin and two additional small molecule inhibitors reduced the viability of YAP1-MAML2-driven mouse tumors *ex vivo*. It remains to be shown if this type of therapy would also be effective in human patients with either *YAP1-MAML2*-positive *NF2*-mutant meningiomas and further investigation will be necessary.

Meningioma is composed of 13 histopathological subtypes with a large histomorphologic spectrum, not including atypical and anaplastic subtypes, which are associated with additional histopathological features. The YAP1-MAML2-driven mouse tumors exhibited histopathological features that are well-described in the spectrum of human meningioma subtypes, including the presence of syncytial cytoplasm (general meningioma), fascicular architecture (fibrous-type), and lobules (meningothelial-type). Histologically, these mouse tumors resemble higher-grade (WHO grade 2 and 3) tumors, likely due to the additional loss of *Cdkn2a* in these tumors. Additional atypical and/or anaplastic histological features present in some of these mouse tumors include areas of patternless growth, macro-nucleoli, pleomorphic nuclei, and elevated mitotic activity, meningioangiomatosis-like growth/invasion. In addition, the YAP1-MAML2- and NLS-2SA-YAP1-driven mouse tumors resembled human YAP1-fusion-driven and NF2-mutant meningiomas in their global and YAP-related gene expression patterns. However, we do recognize that these mouse tumors do not possess all features of classical meningioma (such as prominent whorls). Previously, the Kalamarides lab generated an arachnoid cellspecific PGDS/tv-a mouse line for the modeling of meningioma-like tumors in mice (Peyre et al. 2015). It will be interesting to test if the expression of YAP1-MAML2 or constitutively-active non-fusion YAP1 (S127/397A-YAP1) is able to induce the formation of similar tumors in this the PGDS/tv-a mouse line and if the histomorphology of these tumors will be similar to our tumors.

In summary, our results show that human *NF2*-mutant meningiomas and YAP1-fusion-positive meningiomas express similar gene expression profiles and both harbor enhanced YAP activity. We show that YAP1-MAML2 is a strong oncogenic driver when expressed in mice and the likely tumor-initiating event in *YAP1-MAML2*-positive tumors. The oncogenic functions of YAP1-MAML2 primarily rely on its ability to exert de-regulated TEAD-dependent YAP activity, indicating that YAP1-MAML2 represents an alternative path to achieving de-regulated oncogenic YAP activity in addition to the more common *NF2* loss found in a large percentage of meningiomas. This suggestion is further supported by the similar oncogenic capabilities of constitutively active non-fusion NLS-2SA-YAP1 to also induce the formation of similar tumors, that also resemble human *NF2*-mutant meningiomas by histology and gene expression. Both tumor types similarly responded to pharmacological YAP-TEAD inhibition, suggesting that the YAP component of YAP1-MAML2 is both necessary and sufficient for meningioma formation. This data also indicates that *NF2*-mutant meningiomas may well be dependent on continuous elevated YAP signaling.

## Material and Methods

### Generation of RCAS mouse tumors

All animal experiments were done in accordance with protocols approved by the Institutional Animal Care and Use Committees of Fred Hutchinson Cancer Research Center (FHCRC) (protocol number 50842) and followed NIH guidelines for animal welfare. The RCAS/tv-a system used in this work has been described previously (Szulzewsky et al. 2020). Nestin (N)/tv-a;Cdkn2a null mice were used for RCAS-mediated brain tumor formation in this study and have been described previously (Szulzewsky et al. 2020). 1 × 10^5^ DF1 cells in a volume of 1μl were injected into newborn pup brains, either near the ventricles or into the subarachnoid space (within 1 days after birth). The mice were monitored until they developed symptoms of disease, such as visible tumors, lethargy, poor grooming, weight loss, dehydration, macrocephaly, seizures, jumping, or paralysis, or until a pre-determined study end-point.

### Tissue slice preparation and drug treatments

Tumor slices were prepared as described previously (Sivakumar et al. 2019; Nishida-Aoki et al. 2020). Briefly, dissected tumor tissues were cut into 400 um organotypic tumor slices using the Leica VT1200S vibratome microtome (Nusslock) with HBSS as the cutting medium. The slices were then cut into 400 μm cuboids using a McIlwain Tissue Chopper (Ted Pella) as described previously (Horowitz et al. 2021). Cuboids were immediately placed into 96-well ultra-low attachment plates (Corning) and incubated with Williams’ Medium containing 12 mM nicotinamide, 150 nM ascorbic acid, 2.25 mg/ml sodium bicarbonate, 20 mM HEPES, 50 mg/ml of additional glucose, 1 mM sodium pyruvate, 2 mM L-glutamine, 1% (v/v) ITS, 20 ng/ml EGF, 40 IU/ml penicillin and 40 ug/ml streptomycin containing RealTime Glo reagent (Promega) according to manufacturer’s instructions. After 48 hours, baseline cell viability of cuboids was measured by RealTime Glo bioluminescence using the Synergy H4 instrument (Biotek). Cuboids were exposed to either DMSO (control), Staurosporine (200 nM), Verteporfin or Vivace Therapeutics compounds VT104 and VT107 and overall tumor tissue viability was measured daily, up to 7 days post-treatment.

See also supplemental experimental procedures.

## Supporting information

Suppl. Informtation

## Author Contributions

Conceptualization FSz, PJC, TSG, ECH. Performed experiments FSz, AA. Data analysis FSz, SA, AA, DAB, PS, PJP, FSa, TSG, PJC. Original manuscript writing FSz, SA, PJC, TSG, ECH. Review and editing FSz, AA, PJC, TSG, ECH. Funding acquisition FSz, AA, TSG, ECH. Supervision TSG, ECH. All authors read, reviewed, and approved the manuscript.

## Acknowledgements

We thank Deby Kumasaka, Zachary Russell, Maddie West, and Denis Adair for continued technical and administrative assistance and support throughout these experiments. We thank Alyssa Dawson and Elizabeth Jensen at the Fred Hutchinson Genomics Core for help with DNA sequencing and RNA-Seq. We thank Brianna Wrightson and Elena Carlson for performing the MRI scans on tumor-bearing mice.

## Data and materials availability

The data that support the findings of this study are included with the manuscript and supplemental data files and are also available from the corresponding author upon reasonable request.

## Supplemental Data

Supplemental Figures S1-S7, Supplemental methods

Supplemental Table S1: Sample info human and mouse tumors. Includes data on Sample name, dataset, age, gender, WHO grade, NF2/Chr22 status, TRAF7/KLF4/SMO1/AKT status, and PMID.

Supplemental Table S2: DEGs U5 human neural stem cells expressing S127/397A-YAP1 versus GFP.

Supplemental Table S3: List of reagents used. A) Primers, B) Antibodies, C) siRNA, D) Plasmids.

## References

Antonescu CR, Dickson BC, Sung YS, Zhang L, Suurmeijer AJH, Stenzinger A, Mechtersheimer G, Fletcher CDM. 2020. Recurrent YAP1 and MAML2 Gene Rearrangements in Retiform and Composite Hemangioendothelioma. Am J Surg Pathol 44: 1677–1684.

Baia GS, Caballero OL, Orr BA, Lal A, Ho JS, Cowdrey C, Tihan T, Mawrin C, Riggins GJ. 2012. Yes-associated protein 1 is activated and functions as an oncogene in meningiomas. Mol Cancer Res 10: 904–913.

Barresi V, Simbolo M, Fioravanzo A, Piredda ML, Caffo M, Ghimenton C, Pinna G, Longhi M, Nicolato A, Scarpa A. 2021. Molecular Profiling of 22 Primary Atypical Meningiomas Shows the Prognostic Significance of 18q Heterozygous Loss and CDKN2A/B Homozygous Deletion on Recurrence-Free Survival. Cancers (Basel) 13.

Clark VE, Erson-Omay EZ, Serin A, Yin J, Cotney J, Ozduman K, Avsar T, Li J, Murray PB, Henegariu O et al. 2013. Genomic analysis of non-NF2 meningiomas reveals mutations in TRAF7, KLF4, AKT1, and SMO. Science 339: 1077–1080.

Holley RW, Kiernan JA. 1968. “Contact inhibition” of cell division in 3T3 cells. Proc Natl Acad Sci U S A 60: 300–304.

Horowitz LF, Rodriguez AD, Au-Yeung A, Bishop KW, Barner LA, Mishra G, Raman A, Delgado P, Liu JTC, Gujral TS et al. 2021. Microdissected “cuboids” for microfluidic drug testing of intact tissues. Lab Chip 21: 122–142.

Kalamarides M, Niwa-Kawakita M, Leblois H, Abramowski V, Perricaudet M, Janin A, Thomas G, Gutmann DH, Giovannini M. 2002. Nf2 gene inactivation in arachnoidal cells is rate-limiting for meningioma development in the mouse. Genes Dev 16: 1060–1065.

Liu-Chittenden Y, Huang B, Shim JS, Chen Q, Lee SJ, Anders RA, Liu JO, Pan D. 2012. Genetic and pharmacological disruption of the TEAD-YAP complex suppresses the oncogenic activity of YAP. Genes Dev 26: 1300–1305.

Louis DN, Perry A, Wesseling P, Brat DJ, Cree IA, Figarella-Branger D, Hawkins C, Ng HK, Pfister SM, Reifenberger G et al. 2021. The 2021 WHO Classification of Tumors of the Central Nervous System: a summary. Neuro Oncol 23: 1231–1251.

Maas SLN, Stichel D, Hielscher T, Sievers P, Berghoff AS, Schrimpf D, Sill M, Euskirchen P, Blume C, Patel A et al. 2021. Integrated Molecular-Morphologic Meningioma Classification: A Multicenter Retrospective Analysis, Retrospectively and Prospectively Validated. J Clin Oncol 39: 3839–3852.

Nassiri F, Liu J, Patil V, Mamatjan Y, Wang JZ, Hugh-White R, Macklin AM, Khan S, Singh O, Karimi S et al. 2021. A clinically applicable integrative molecular classification of meningiomas. Nature 597: 119–125.

Nishida-Aoki N, Bondesson AJ, Gujral TS. 2020. Measuring Real-time Drug Response in Organotypic Tumor Tissue Slices. J Vis Exp.

O’Connor SA, Feldman HM, Arora S, Hoellerbauer P, Toledo CM, Corrin P, Carter L, Kufeld M, Bolouri H, Basom R et al. 2021. Neural G0: a quiescent-like state found in neuroepithelial-derived cells and glioma. Mol Syst Biol 17: e9522.

Ostrom QT, Patil N, Cioffi G, Waite K, Kruchko C, Barnholtz-Sloan JS. 2020. CBTRUS Statistical Report: Primary Brain and Other Central Nervous System Tumors Diagnosed in the United States in 2013-2017. Neuro Oncol 22: iv1–iv96.

Ozawa T, Arora S, Szulzewsky F, Juric-Sekhar G, Miyajima Y, Bolouri H, Yasui Y, Barber J, Kupp R, Dalton J et al. 2018. A De Novo Mouse Model of C11orf95-RELA Fusion-Driven Ependymoma Identifies Driver Functions in Addition to NF-kappaB. Cell Rep 23: 3787–3797.

Pajtler KW, Wei Y, Okonechnikov K, Silva PBG, Vouri M, Zhang L, Brabetz S, Sieber L, Gulley M, Mauermann M et al. 2019. YAP1 subgroup supratentorial ependymoma requires TEAD and nuclear factor I-mediated transcriptional programmes for tumorigenesis. Nat Commun 10: 3914.

Patel AJ, Wan YW, Al-Ouran R, Revelli JP, Cardenas MF, Oneissi M, Xi L, Jalali A, Magnotti JF, Muzny DM et al. 2019. Molecular profiling predicts meningioma recurrence and reveals loss of DREAM complex repression in aggressive tumors. Proc Natl Acad Sci U S A 116: 21715–21726.

Petrilli AM, Fernandez-Valle C. 2016. Role of Merlin/NF2 inactivation in tumor biology. Oncogene 35: 537–548.

Peyre M, Salaud C, Clermont-Taranchon E, Niwa-Kawakita M, Goutagny S, Mawrin C, Giovannini M, Kalamarides M. 2015. PDGF activation in PGDS-positive arachnoid cells induces meningioma formation in mice promoting tumor progression in combination with Nf2 and Cdkn2ab loss. Oncotarget 6: 32713–32722.

Peyre M, Stemmer-Rachamimov A, Clermont-Taranchon E, Quentin S, El-Taraya N, Walczak C, Volk A, Niwa-Kawakita M, Karboul N, Giovannini M et al. 2013. Meningioma progression in mice triggered by Nf2 and Cdkn2ab inactivation. Oncogene 32: 4264–4272.

Pobbati AV, Hong W. 2020. A combat with the YAP/TAZ-TEAD oncoproteins for cancer therapy. Theranostics 10: 3622–3635.

Prager BC, Vasudevan HN, Dixit D, Bernatchez JA, Wu Q, Wallace LC, Bhargava S, Lee D, King BH, Morton AR et al. 2020. The Meningioma Enhancer Landscape Delineates Novel Subgroups and Drives Druggable Dependencies. Cancer Discov 10: 1722–1741.

Riemenschneider MJ, Perry A, Reifenberger G. 2006. Histological classification and molecular genetics of meningiomas. Lancet Neurol 5: 1045–1054.

Sahm F, Schrimpf D, Stichel D, Jones DTW, Hielscher T, Schefzyk S, Okonechnikov K, Koelsche C, Reuss DE, Capper D et al. 2017. DNA methylation-based classification and grading system for meningioma: a multicentre, retrospective analysis. Lancet Oncol 18: 682–694.

Sekine S, Kiyono T, Ryo E, Ogawa R, Wakai S, Ichikawa H, Suzuki K, Arai S, Tsuta K, Ishida M et al. 2019. Recurrent YAP1-MAML2 and YAP1-NUTM1 fusions in poroma and porocarcinoma. J Clin Invest 129: 3827–3832.

Sievers P, Chiang J, Schrimpf D, Stichel D, Paramasivam N, Sill M, Gayden T, Casalini B, Reuss DE, Dalton J et al. 2020a. YAP1-fusions in pediatric NF2-wildtype meningioma. Acta Neuropathol 139: 215–218.

Sievers P, Hielscher T, Schrimpf D, Stichel D, Reuss DE, Berghoff AS, Neidert MC, Wirsching HG, Mawrin C, Ketter R et al. 2020b. CDKN2A/B homozygous deletion is associated with early recurrence in meningiomas. Acta Neuropathol 140: 409–413.

Sivakumar R, Chan M, Shin JS, Nishida-Aoki N, Kenerson HL, Elemento O, Beltran H, Yeung R, Gujral TS. 2019. Organotypic tumor slice cultures provide a versatile platform for immuno-oncology and drug discovery. Oncoimmunology 8: e1670019.

Stein C, Bardet AF, Roma G, Bergling S, Clay I, Ruchti A, Agarinis C, Schmelzle T, Bouwmeester T, Schubeler D et al. 2015. YAP1 Exerts Its Transcriptional Control via TEAD-Mediated Activation of Enhancers. PLoS Genet 11:e1005465.

Striedinger K, VandenBerg SR, Baia GS, McDermott MW, Gutmann DH, Lal A. 2008. The neurofibromatosis 2 tumor suppressor gene product, merlin, regulates human meningioma cell growth by signaling through YAP. Neoplasia 10: 1204–1212.

Szulzewsky F, Arora S, Hoellerbauer P, King C, Nathan E, Chan M, Cimino PJ, Ozawa T, Kawauchi D, Pajtler KW et al. 2020. Comparison of tumor-associated YAP1 fusions identifies a recurrent set of functions critical for oncogenesis. Genes Dev 34: 1051–1064.

Szulzewsky F, Holland EC, Vasioukhin V. 2021. YAP1 and its fusion proteins in cancer initiation, progression and therapeutic resistance. Dev Biol 475: 205–221.

Tanahashi K, Natsume A, Ohka F, Motomura K, Alim A, Tanaka I, Senga T, Harada I, Fukuyama R, Sumiyoshi N et al. 2015. Activation of Yes-Associated Protein in Low-Grade Meningiomas Is Regulated by Merlin, Cell Density, and Extracellular Matrix Stiffness. J Neuropathol Exp Neurol 74: 704–709.

Tang TT, Konradi AW, Feng Y, Peng X, Ma M, Li J, Yu FX, Guan KL, Post L. 2021. Small Molecule Inhibitors of TEAD Auto-palmitoylation Selectively Inhibit Proliferation and Tumor Growth of NF2-deficient Mesothelioma. Mol Cancer Ther 20: 986–998.

Zhang N, Bai H, David KK, Dong J, Zheng Y, Cai J, Giovannini M, Liu P, Anders RA, Pan D. 2010. The Merlin/NF2 tumor suppressor functions through the YAP oncoprotein to regulate tissue homeostasis in mammals. Dev Cell 19: 27–38.

Zhao B, Li L, Tumaneng K, Wang CY, Guan KL. 2010. A coordinated phosphorylation by Lats and CK1 regulates YAP stability through SCF(beta-TRCP). Genes Dev 24: 72–85.

Zhao B, Wei X, Li W, Udan RS, Yang Q, Kim J, Xie J, Ikenoue T, Yu J, Li L et al. 2007. Inactivation of YAP oncoprotein by the Hippo pathway is involved in cell contact inhibition and tissue growth control. Genes Dev 21: 2747–2761.

Zhao B, Ye X, Yu J, Li L, Li W, Li S, Yu J, Lin JD, Wang CY, Chinnaiyan AM et al. 2008. TEAD mediates YAP-dependent gene induction and growth control. Genes Dev 22: 1962–1971.

