## Supplementary material for "Both YAP1-MAML2 and constitutively active YAP1 drive the formation of tumors that resemble NF2-mutant meningiomas in mice": Suppl. Informtation

Supplemental Information

**Suppl. Figures:**

**Suppl. Figure S1: Human NF2-wild type YAP1 fusion-positive meningiomas resemble NF2-mutant meningiomas by gene expression.** A) UMAP of human meningioma and PA samples. Samples are colored in by dataset, mutation status, methylation subtype, or gender. The UMAP was calculated based on all samples in the dataset (including mouse tumor samples), but only human samples are shown. B) UMAP of human meningioma and PA samples. Samples are colored in by WHO grade, NF2mutation/Chr22 status, mutation status, or methylation subtype. The UMAP was calculated only with samples shown on the plot (excluding mouse tumor samples). C) Hierarchical clustering of human meningioma and PA samples. Samples are colored in by TRAF7/KLF4/AKT1/SMO1 mutation status, methylation subtype, NF2mutation/Chr22 status, WHO grade, and dataset.

**Suppl. Figure S2: Human YAP1 fusion-positive meningiomas and NF2-mutant meningiomas exert increased levels of YAP signaling.** A) Expression of selected YAP1 target genes (*AJUBA, ANXA1, ANXA3, and CITED2*) in human meningiomas and PAs. B-C) Clustering of human samples based on the expression of 2SA-YAP1 up- (B) and down-regulated (C) genes. Analysis was done using ordinary one-way ANOVA (A). (*) $P < 0.05$; (**) $P < 0.01$.

**Suppl. Figure S3: Forced expression of YAP1-MAML2 induces the formation of meningioma-like tumors in mice.** A) Schematic of wtYAP1, wtMAML2, full length YAP1(e1)-MAML2 (YM), YM-v1, YM-v2, truncated wtMAML2-v1 and -v2, and full length YAP1(e1-e5)-MAML2 (YM) protein sequences. Amino acid positions shown are excluding the HA-tag sequence (9 amino acids). WtYAP1, YM-v1, YM-v2, truncated wtMAML2-v1 and truncated wtMAML2-v2 were used for in vivo experiments. B) Western Blot showing expression of HA-tagged versions of wtYAP1, NLS-2SA-YAP1, YM-v1, YM-v2, truncated wtMAML2-v1, truncated wtMAML2-v2, and S94A-YM-v2 in DF1 cells. Blot was stained with anti-HA and anti-Actin antibodies. C) YAP activity of GFP, wtYAP1, and full length YM, YM-v1, and YM-v2 in the YAP1-responsive GTIIC-Luc reporter assay (n = 2). D) Nestin immunohistochemistry (IHC) staining of a p0 neonatal mouse brain and skull. Nestin staining was observed both in the meninges (arrowhead) and the sub-ventricular zone (arrow). E) H&E and HA stainings of small lesions near the ventricles induced by the intra-cranial expression of YM-v2 in N/tva Cdkn2a wild type mice. F) Injection summary for YM-v2 in N/tv-a Cdkn2a wild type mice. G) H&E staining of an intra-ventricular tumor induced by expression of YM-v2 in N/tv-a Cdkn2a null mice. H) Injection summaries for YM-v1, wtMAML2-v1, wtMAML2-v2, and wtYAP1 in N/tv-a Cdkn2a null mice. I) Upper panel: T2w MRI images taken 77, 90, or 99 days post-injection. Lower panel: T1w MRI images pre- (left) or post-contrast (middle) taken 102 days post-injection. H&E staining of the same tumor (right). J-J) Higher power images of H&E stainings of representative YM-v1 and YM-v2 tumors. S) Focal EMA IHC staining in a representative YM-v2 tumor. T) MUC1 (EMA) expression in mouseYM samples, human meningiomas, and PA samples. U) IHC stainings for HA (tumor cells), Vimentin, Synaptophysin, GFAP, and OLIG2 of a representative YM-v2 extra-axial tumor. The tumor-brain surface is shown, the tumor is located on the upper left side. V) IHC stainings for OLIG2, showing an area devoid of

OLIG2-positive cells (left) and an area with several OLIG2-positive cells. W) Co-immunofluorescence stainings for HA (tumor cells) and OLIG2 showing no overlap between the two markers, suggesting that the OLIG2-positive cells are entrapped untransformed-glia cells. X) UMAP of mouseYM samples, human meningioma and PA samples. Samples are colored in by WHO grade, NF2mutation/Chr22 status, mutation status, or methylation subtype. The UMAP was calculated based on all samples in the dataset (including mouse2SA tumor samples), but only mouseYM and human samples are shown. Y) UMAP of mouseYM samples, human meningioma and PA samples. Samples are colored in by WHO grade, NF2mutation/Chr22 status, mutation status, or methylation subtype. The UMAP was calculated only with samples shown on the plot (excluding mouse2SA tumor samples). Z-AA) Clustering of mouseYM and human samples based on the expression of 2SA-YAP1 up- (Z) and down-regulated (AA) genes. AB) Expression of selected YAP1 target genes (*CTGF, CYR61, AMOTL2, ANKRD1, CPA4, AJUBA, ANXA1, ANXA3, and CITED2*) in mouseYM tumors, human meningiomas, and PAs. Error bars show SEM. Analysis was done using ordinary one-way ANOVA (C,S,X). (*) $P < 0.05$; (**) $P < 0.01$; (***) $P < 0.001$; (****) $P < 0.0001$.

**Suppl. Figure S4: YAP1-MAML2 is constitutively localized to the nucleus and is insensitive to Hippo pathway-mediated inhibition.** A) HA IHC staining of a YM-v1 mouse tumor demonstrates nuclear localization of YM in vivo. B) Western Blot showing expression of HA-tagged versions of wtYAP1, S94A-YAP1, YAP1(e1)-MAML2(e2-e5) full length, YAP1(e1-e5)-MAML2(e2-e5) full length, S94A-YAP1(e1)-MAML2(e2-e5) full length, YM-v1, YM-v2, and S94A-YM-v2 in DF1 cells. Blot was stained with anti-HA and anti-Actin antibodies.

**Suppl. Figure S5: The transcriptional program induced by YAP1-MAML2 is dependent on the interaction with TEAD transcription factors.** A) Expression of the YAP1 target genes *CPA4*, *AJUBA*, *ANXA1*, and *ANXA3* in GFP-, wtYAP1-, YM-, or S94A-YM-expressing HEK cells from RNA-Seq data. B) Volcano plot showing DEGs induced in YM-expressing HEK cells compared to GFP-expressing cells. C) Volcano plot showing DEGs induced in wtYAP1-expressing HEK cells compared to GFP-expressing cells. D) Volcano plot showing DEGs induced in S94A-YM-expressing HEK cells compared to GFP-expressing cells. E) Volcano plot showing the regulation of YM-DEGs in S94A-YM-expressing HEK. F) Expression of TEAD1-4 in mouseYM tumors, human meningiomas, and PAs. G) TEAD1 IHC staining in naïve brain and YM-v1 mouse tumors. H) YAP activity of S94A mutant wtYAP1, YM, and YM-v2 in the GTIIC-Luc reporter assay (n = 4 each). Error bars show SEM. Analysis was done using ordinary one-way ANOVA (A,F) or two-tailed t-test (H). (*) $P < 0.05$; (**) $P < 0.01$; (***) $P < 0.001$; (****) $P < 0.0001$.

**Suppl. Figure S6: The interaction with TEAD transcription factors is necessary for the oncogenic activity of YAP1-MAML2 in vivo.** A-C) Spheroid growth of untransduced (A) NIH3T3 cells or cells expressing either wtYAP1 (B) or wtMAML2 (C) when treated with VP or DMSO only (n = 3 each). D) Expression of *Ctgf* and *Cyr61* in YM-expressing NIH3T3 cells treated with either DMSO (control) or Verteporfin. E) Viability of YM-v2 organotypic mouse tumor cuboids (tumor #4) after no treatment (NT) or treatment with either DMSO, Staurosporine (SPS), Verteporfin, or

VT104/VT107. F) Expression of *Ctgf* and *Cyr61* in YM-v2 mouse tumor cuboids treated with either DMSO (control) or Verteporfin for 7 days. Error bars show SD. Analysis was done using ordinary two-way ANOVA (A-C), ordinary one-way ANOVA (E), or two-tailed t-test (D, F). (*) $P < 0.05$; (**) $P < 0.01$; (***) $P < 0.001$; (****) $P < 0.0001$.

**Suppl. Figure S7: Expression of constitutively activated YAP1 itself is sufficient to cause the formation of meningioma-like tumors in mice.** A-B) PCA plot (A) and Hierarchical clustering (B) showing the similarity of RNA-Seq samples of U5 human neural stem cells (wild type (untreated), GFP, wtYAP1, 2SA-YAP1, sgCTL, sgNF2). C) Overlap between the DEGs of sgNF2 and 2SA-YAP1-expressing U5 human neural stem cells (compared to untreated wild type cells). D) Expression of YAP1 and YAP1 target genes in wild type (untreated), sgNF2, or 2SA-YAP1-expressing U5 human neural stem cells. E-S) Representative higher power images of NLS-2SA-YAP1-induced meningioma-like mouse tumors. T-U) Representative IHC staining for the HA-tag (T) and TEAD1 (U) in a NLS-2SA-YAP1 induced extra-axial mouse tumor. V) UMAP of mouse2SA, mouseYM, human meningioma, and PA samples. Samples are colored in by WHO grade, NF2mutation/Chr22 status, mutation status, or methylation subtype. W-X) Clustering of mouse2SA, mouseYM, and human samples based on the expression of 2SA-YAP1 up- (W) and down-regulated (X) genes. Y) Expression of selected YAP1 target genes (*CTGF, CYR61, AMOTL2, ANKRD1, CPA4, AJUBA, ANXA1, ANXA3, and CITED2*) in mouse2SA, mouseYM, human meningioma, and PA samples. Z) Viability of NLS-2SA-YAP1 organotypic mouse tumor cuboids (taken from five separate extra-cranial tumors) after no treatment (NT) or treatment with either DMSO, Staurosporine (STS), Verteporfin, or VT104/VT107. Error bars show SD. Analysis was done using ordinary one-way ANOVA (D,Y,Z). (*) $P < 0.05$; (**) $P < 0.01$; (***) $P < 0.001$; (****) $P < 0.0001$.

Suppl. Figure S1

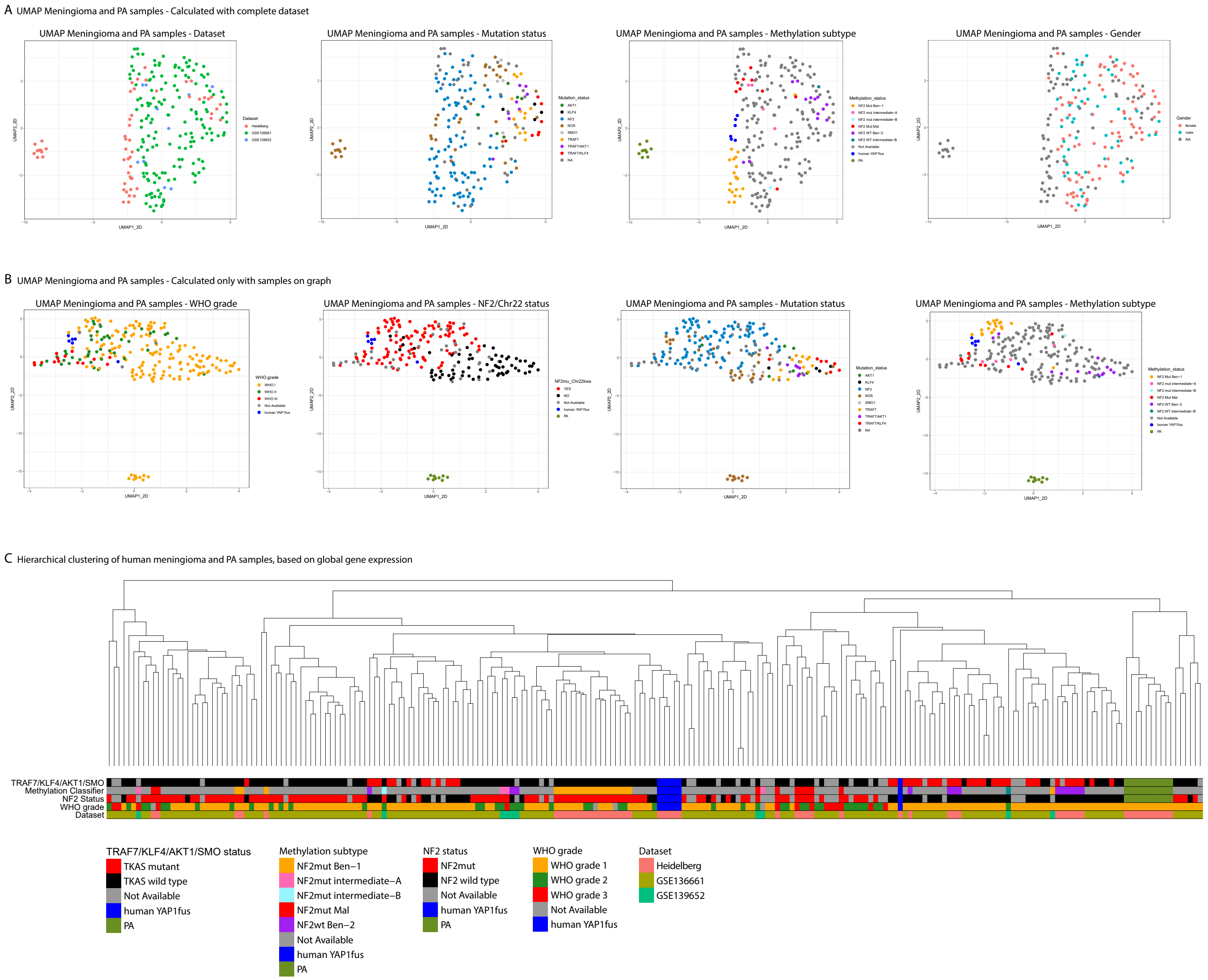

### Suppl. Figure S2

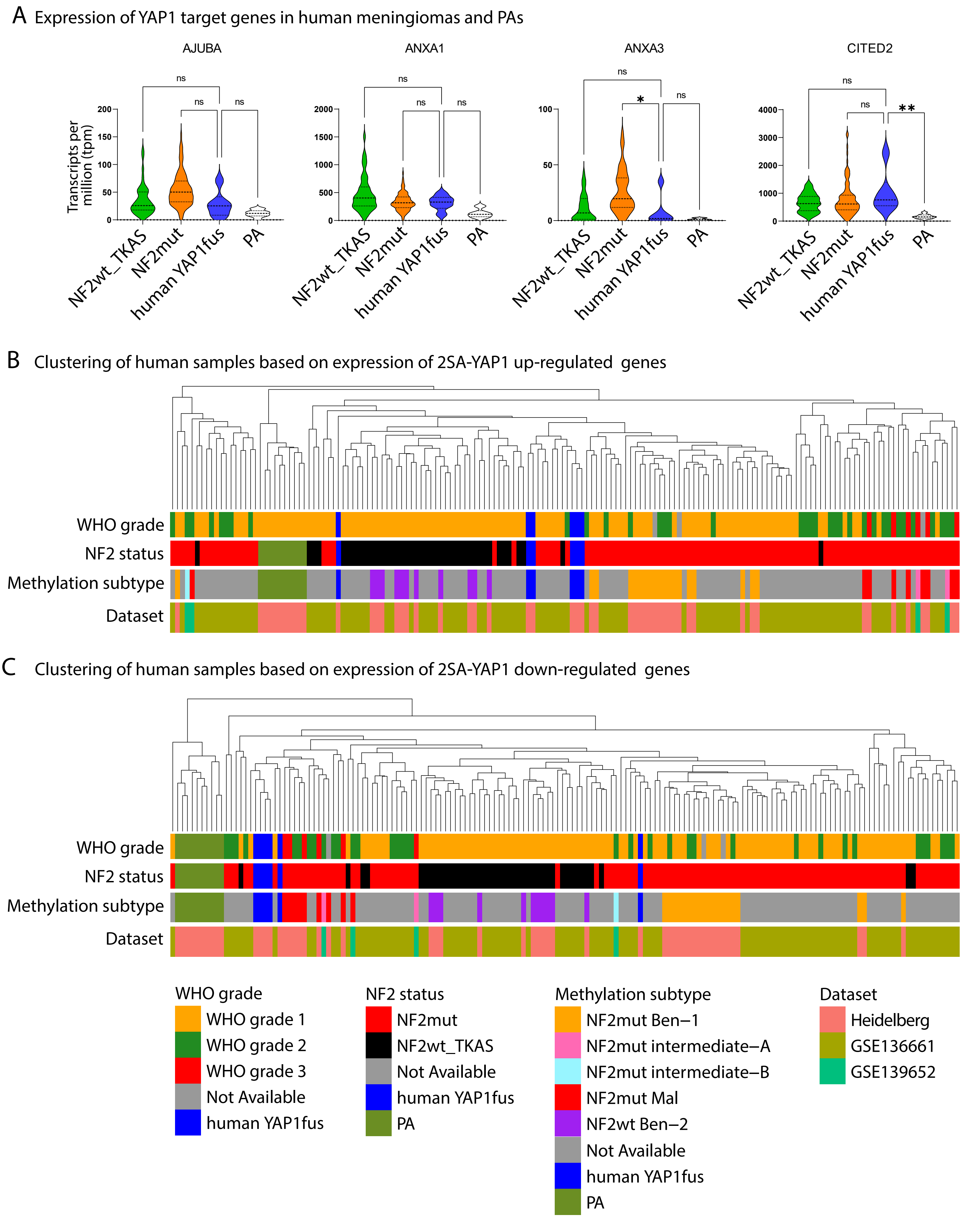

### Suppl. Figure S3A-C

A Schematic of wtYAP1, wtMAML2, full length YAP1(e1)-MAML2, truncated versions YAP1-MAML2-v1 and -v2, truncated wtMAML2-v1 and -v2, and full length YAP1(e1)-MAML2.

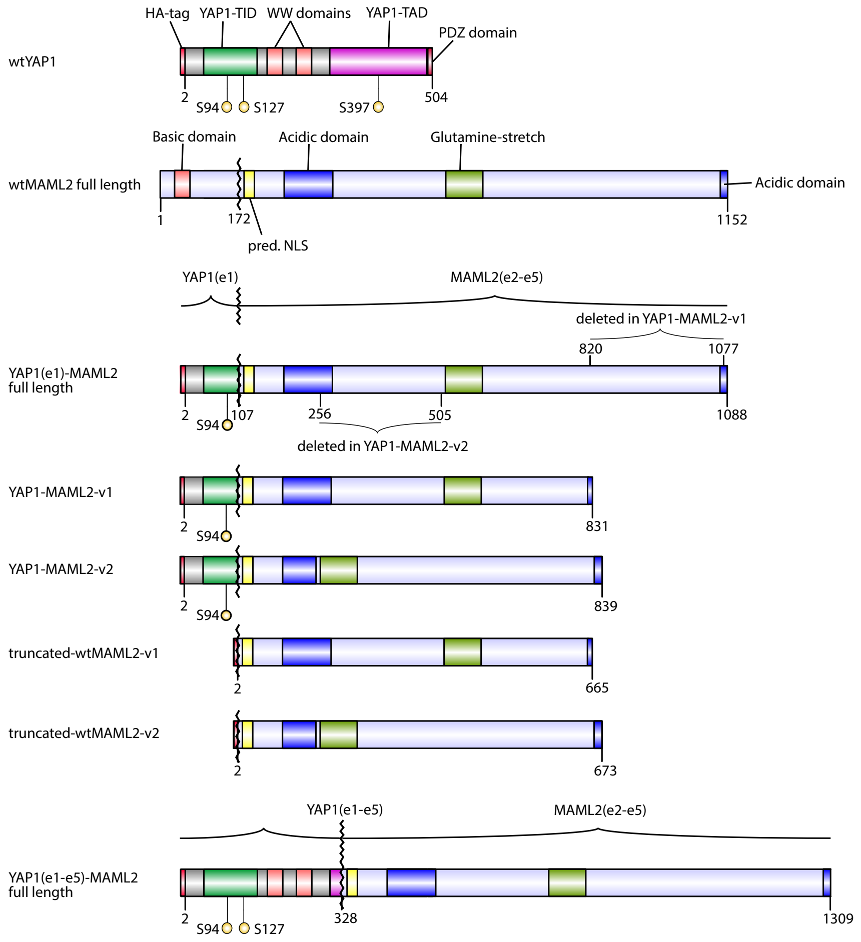


B Western Blot of DF1 cells expressing different RCAS constructs

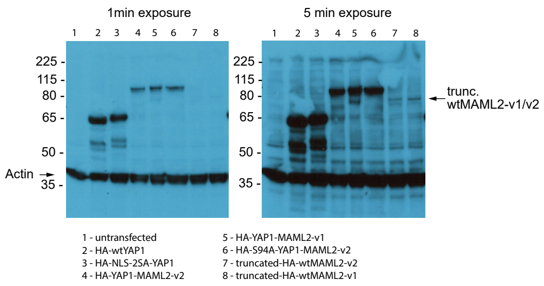


C

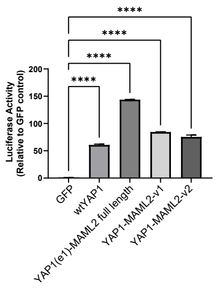

### Suppl. Figure S3D-H

D Nestin/tv-a *Cdkn2a* null p0 neonatal brain

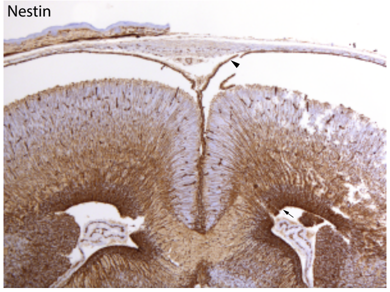


E RCAS-YMv2 intra-ventricular

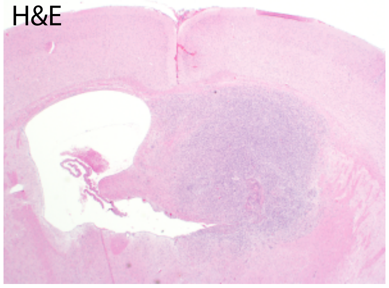


F RCAS-YMv2 intra-ventricular

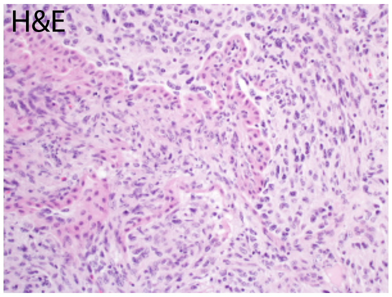


G Injection in Nestin/tv-a Cdkn2a null mice

| **YAP1-MAML2-v1** | No of mice injected | No of mice with tumors | Penetrance (Percent) | Intra-ventricular | Extra-axial | Extracranial | Latency (Days) |
|---|---|---|---|---|---|---|---|
| **DEEP INJECTION** | 12 | 5 | 41.7 | 2 | 1 | 2 | 87-175 |

| | No of mice injected | No of mice with tumors | Penetrance (Percent) | Latency (Days) |
|---|---|---|---|---|
| **WTMAML2-V1** | 5 | 0 | 0 | 150 |
| **WTMAML2-V2** | 15 | 0 | 0 | 150 |
| **WTYAP1** | 8 | 0 | 0 | 200 |

H Days post-injection

77 days 90 days 99 days

T2 weighted

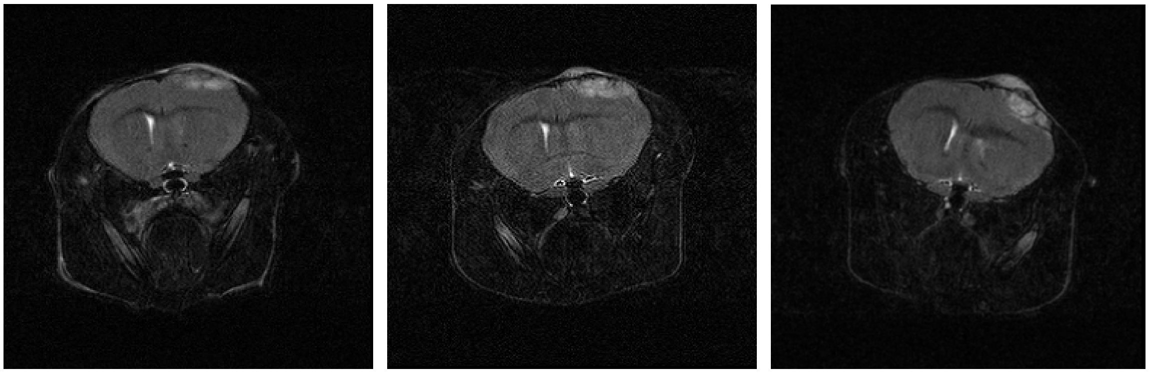

pre-contrast 10 min post-contrast H&E

T1 weighted (102 days post-injection)

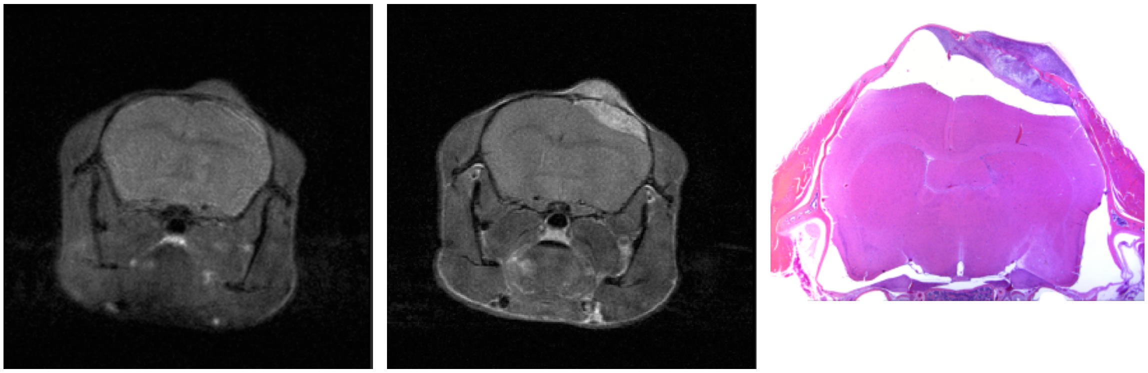

Suppl. Figure S3I

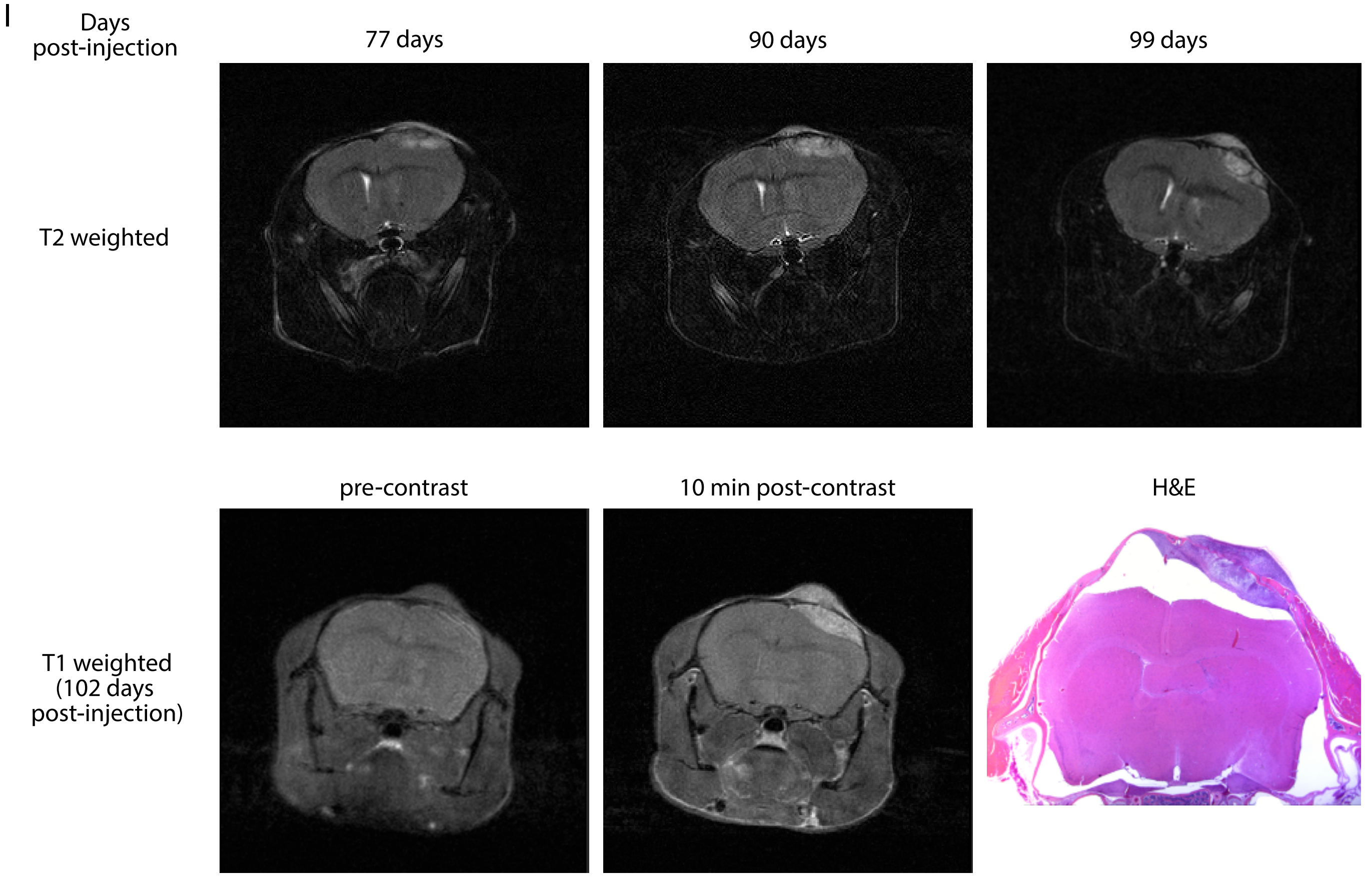

Suppl. Figure S3J-T

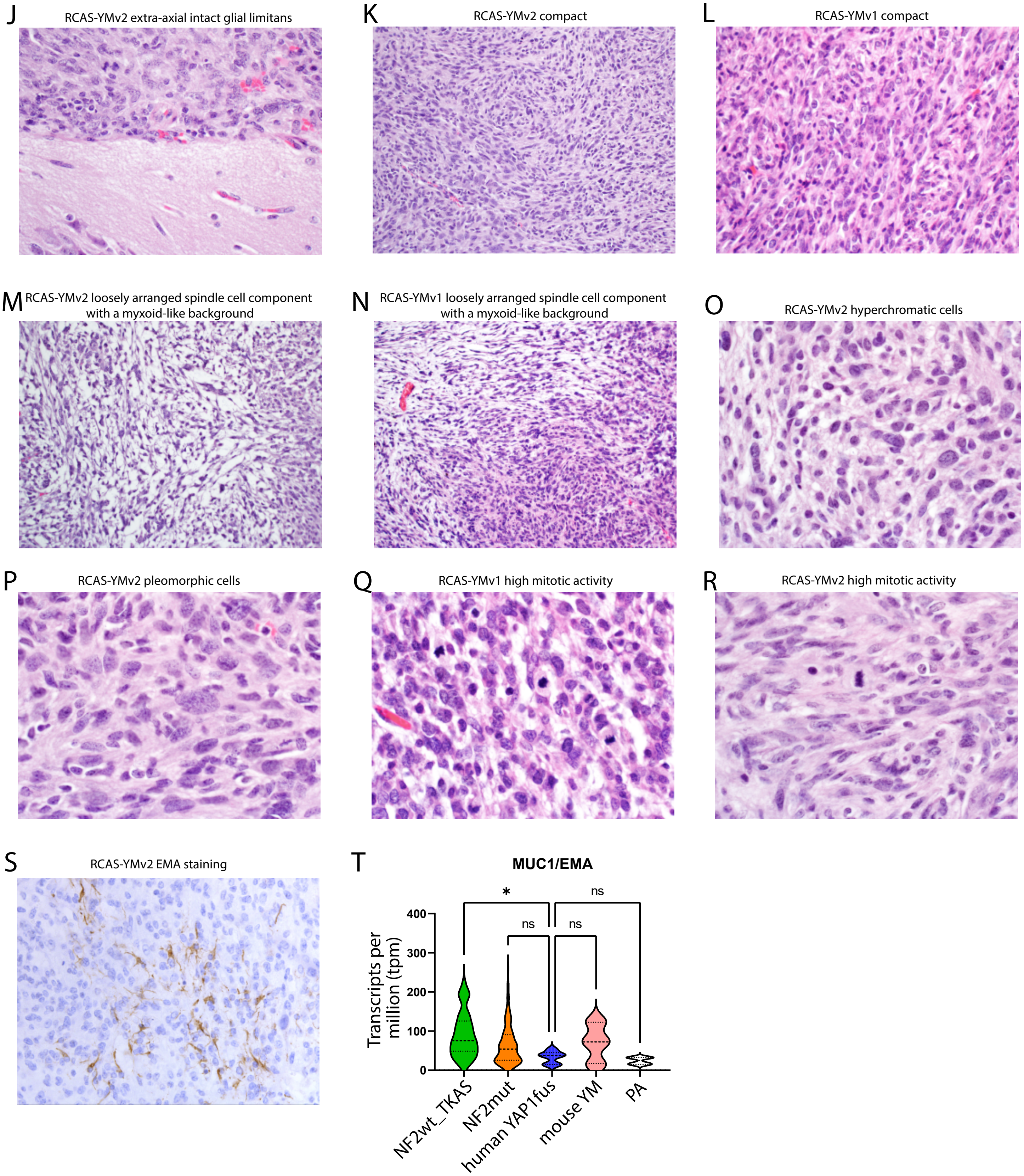

Suppl. Figure S3U-W

U IHC stainings of RCAS-YMv2 tumors

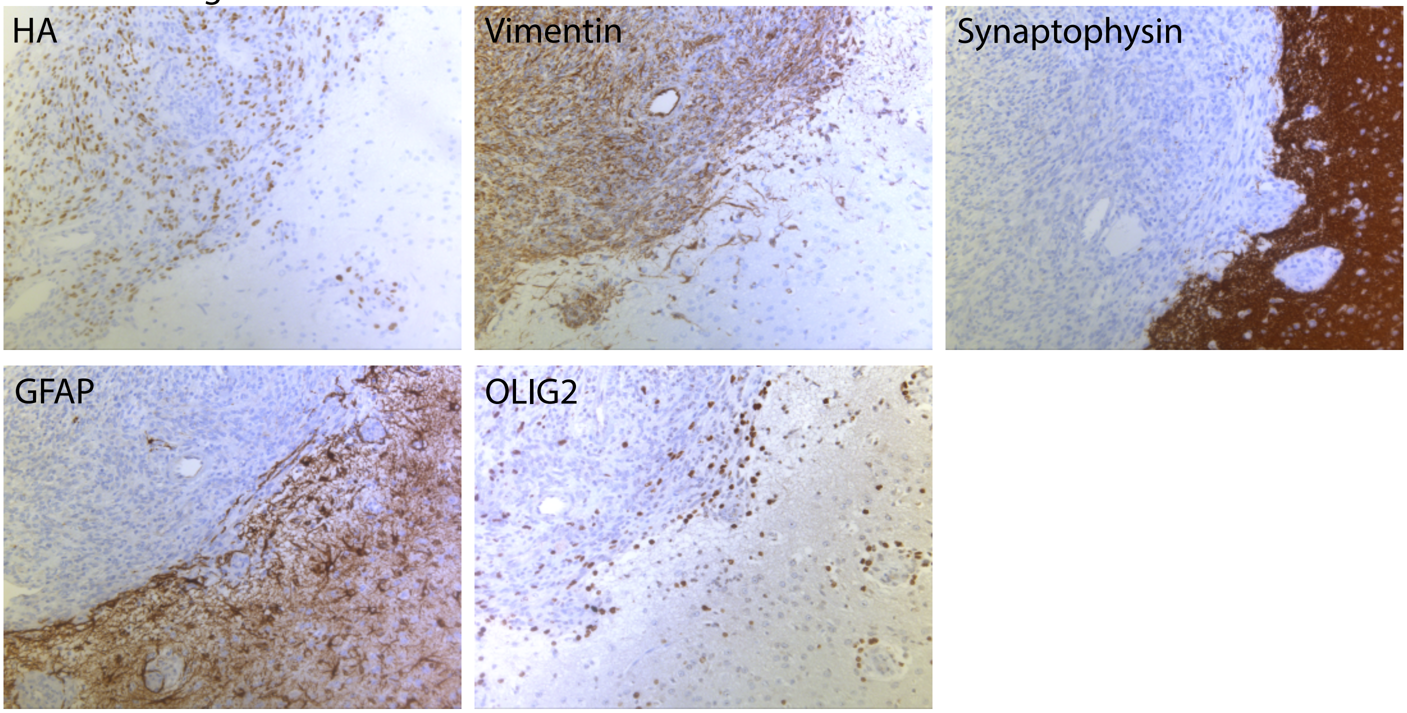


V

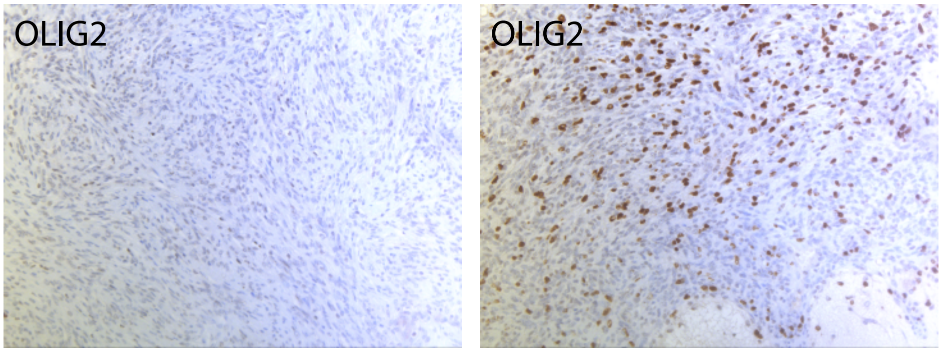


W Co-IF stainings of RCAS-YMv2 tumors for OLIG2 and HA

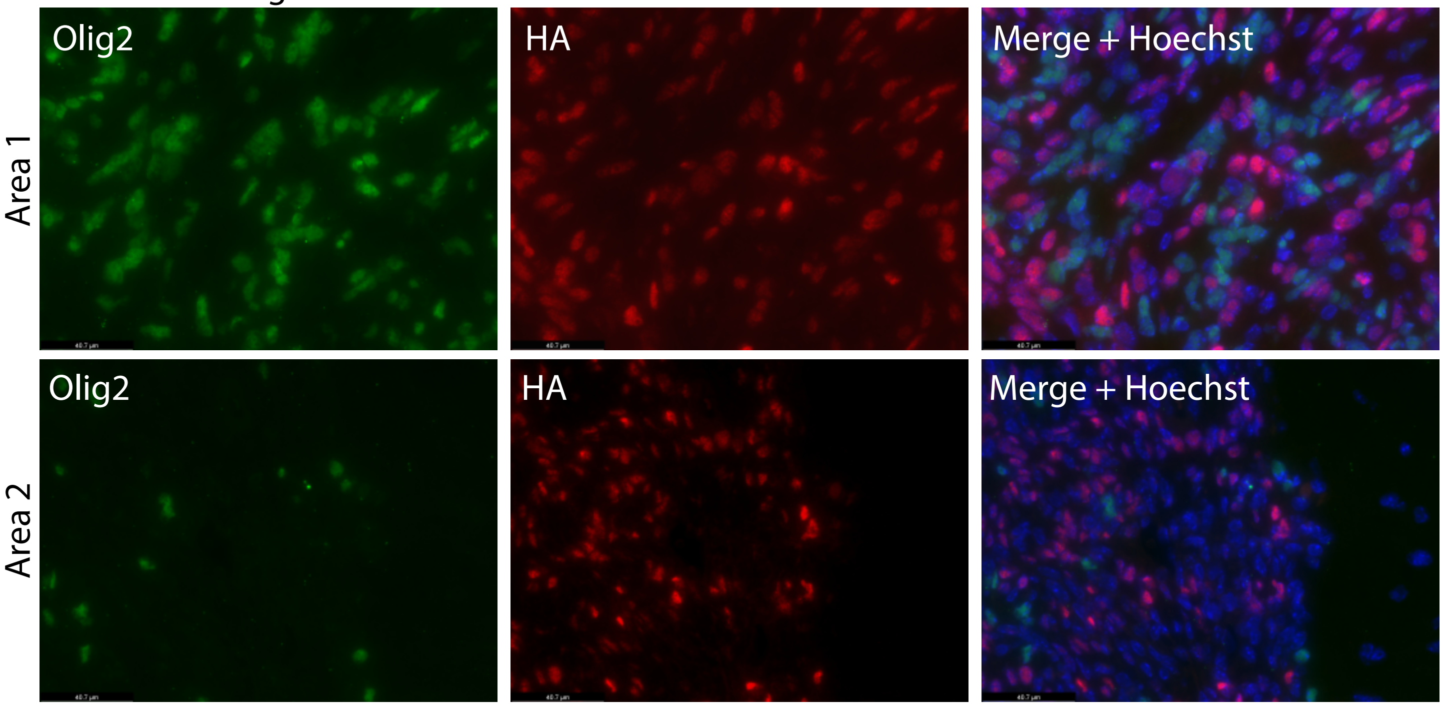

### Suppl. Figure S3X-AB

#### X UMAP Meningioma and PA samples - Calculated with complete dataset

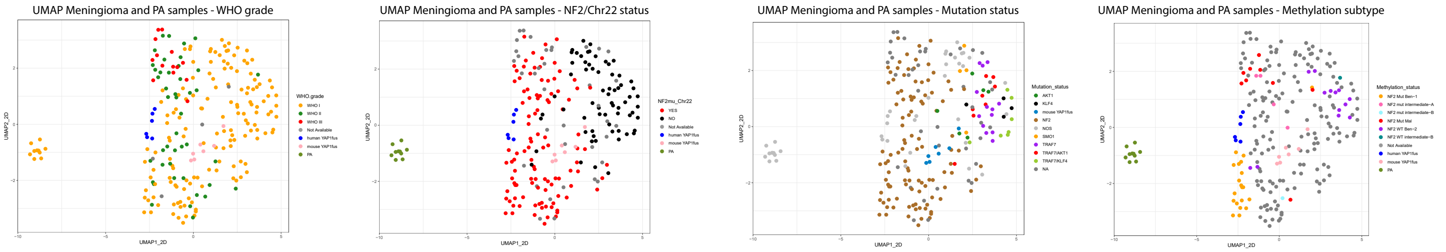


#### Y UMAP Meningioma and PA samples - Calculated only with samples on graph

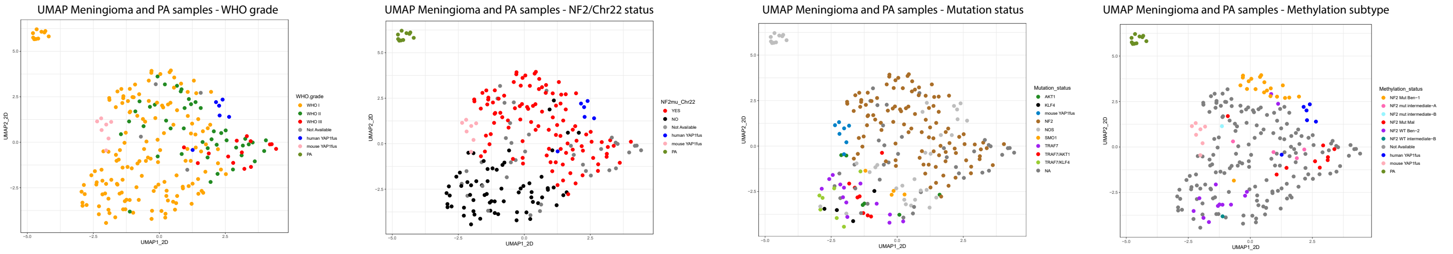


#### Z Clustering of human and mouseYM samples based on expression of 2SA-YAP1 up-regulated genes

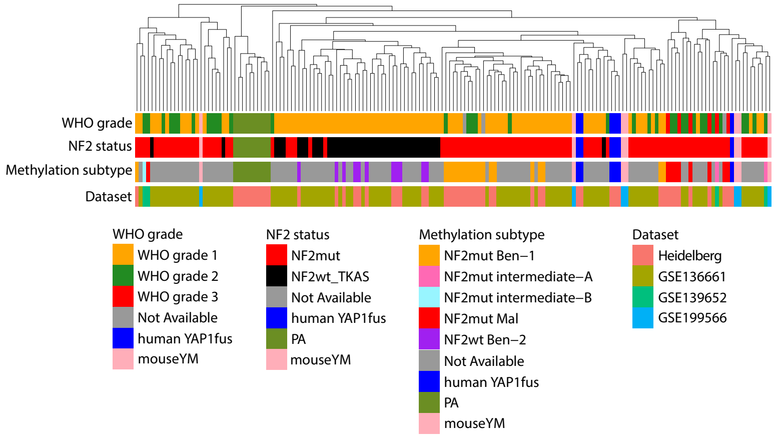


#### AA Clustering of human and mouseYM samples based on expression of 2SA-YAP1 down-regulated genes

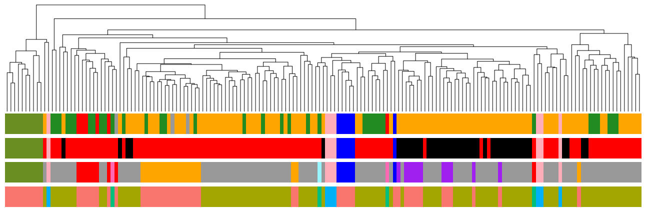

#### AB Expression of YAP1 target genes in human meningiomas, mouse YM tumors, and PAs

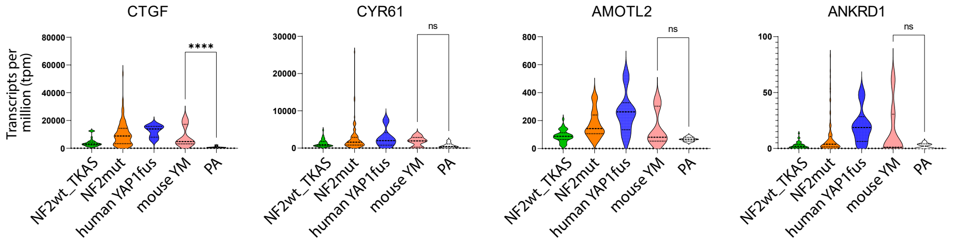


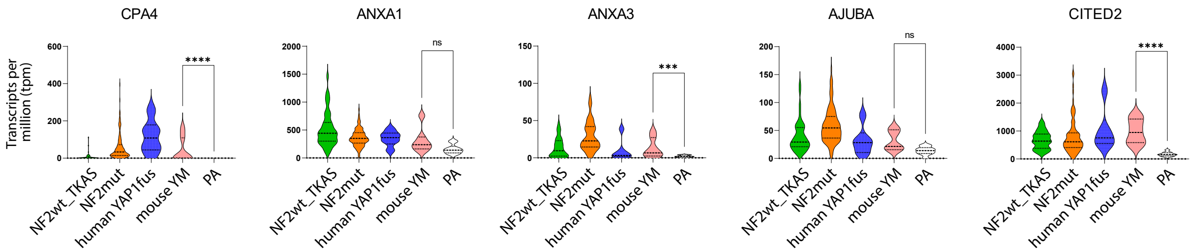

Suppl. Figure S4

A YAP1-MAML2-v1

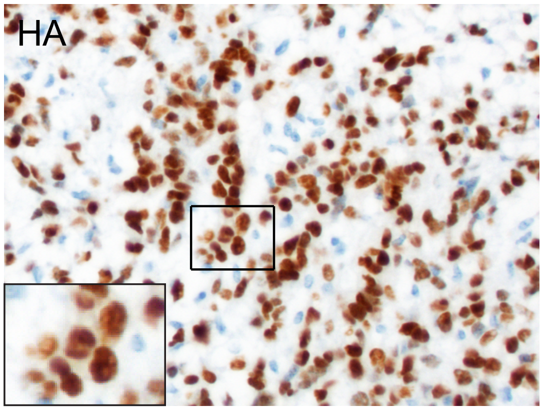


B Western Blot HEK cells expressing different RCAS constructs

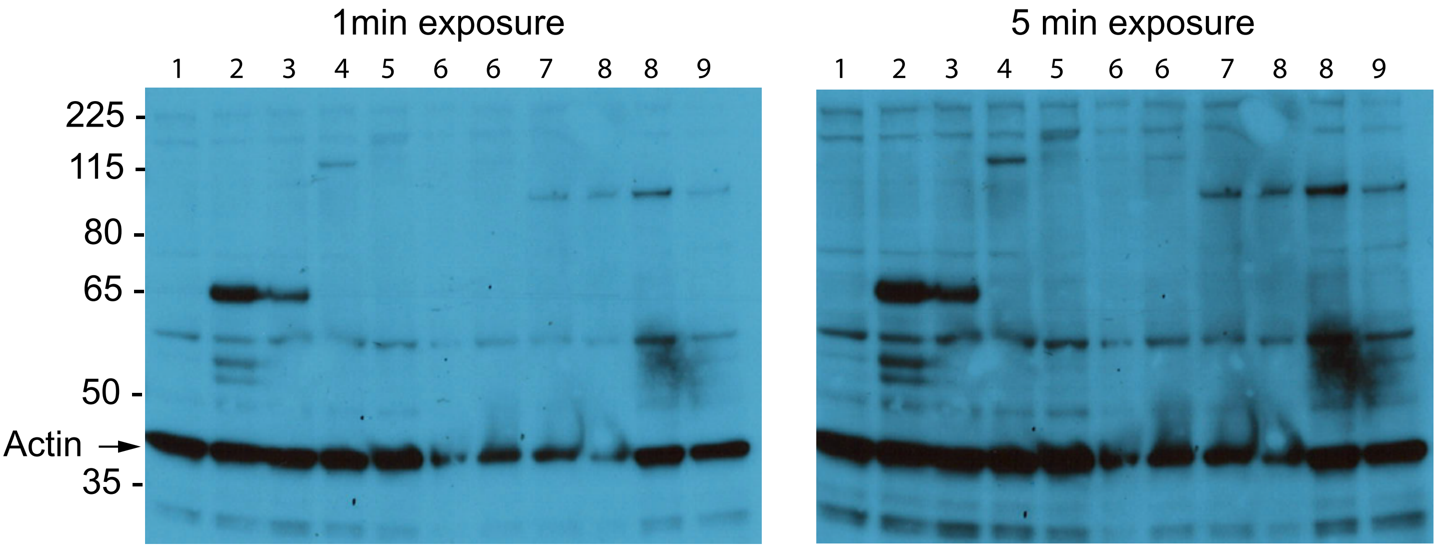


1 - untransfected
2 - HA-wtYAP1
3 - HA-S94A-YAP1
4 - HA-YAP1(e1)-MAML2 full length
5 - HA-YAP1(e1-e5)-MAML2 full length
6 - HA-S94A-YAP1(e1)-MAML2(e2-e5) full length
7 - HA-YAP1-MAML2-v2
8 - HA-YAP1-MAML2-v1
9 - HA-S94A-YAP1-MAML2-v2

Suppl. Figure S5A-E

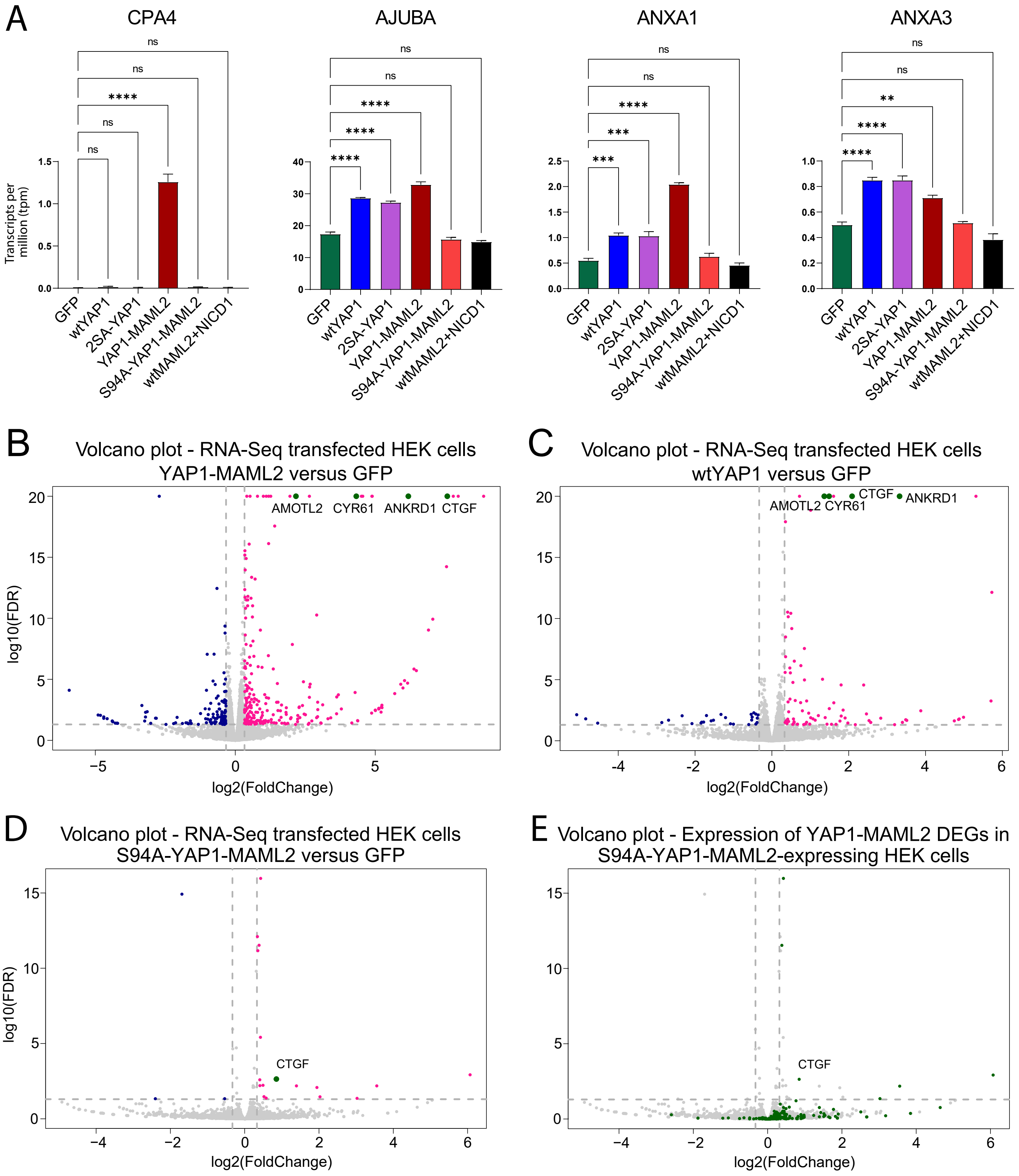

### Suppl. Figure S3F-H

F

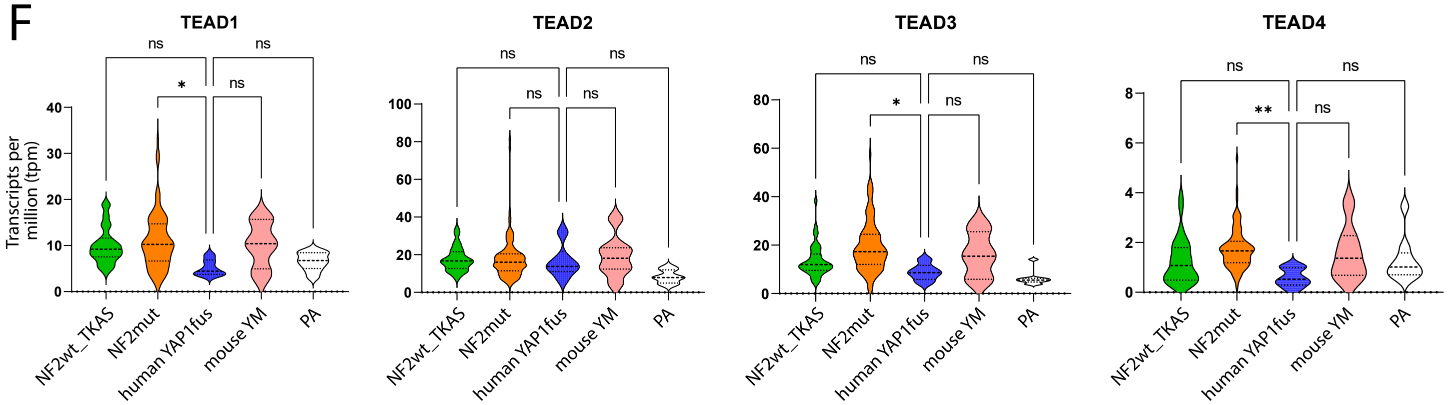


G

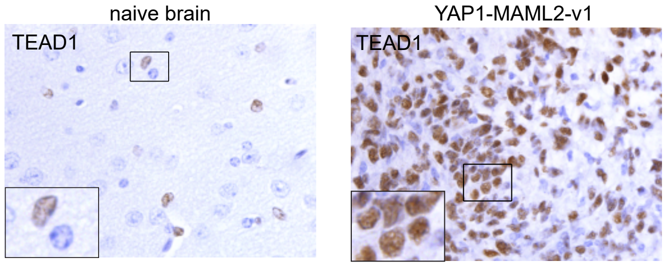


H

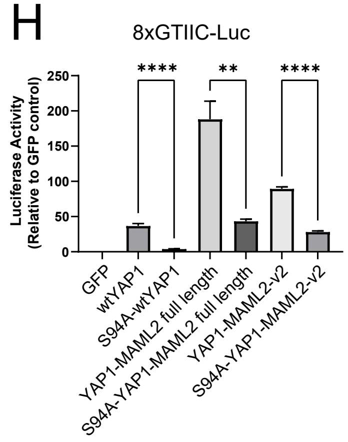

#### Suppl. Figure S6

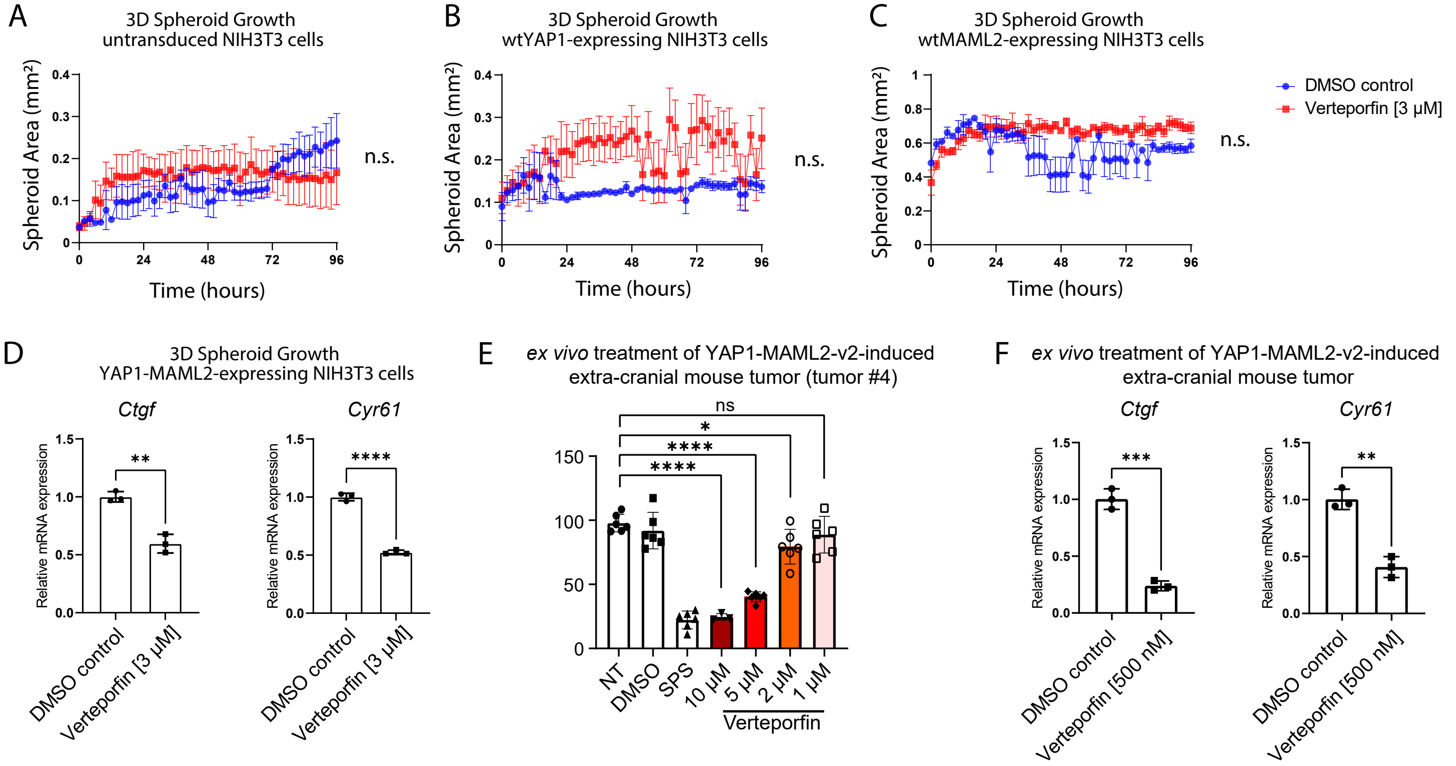

Suppl. Figure S7A-D

A
PCA plot - RNA-Seq U5 human neural stem cells
Comparison 2SA-YAP1 and sgNF2 cells

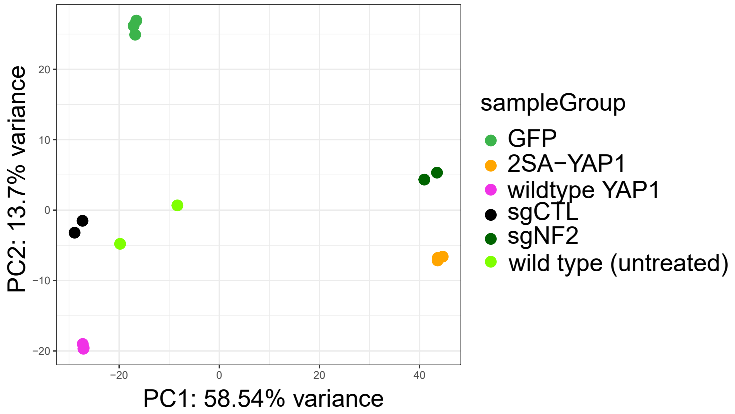


B
Hierarchical clustering - RNA-Seq U5 human neural stem cells
Comparison 2SA-YAP1 and sgNF2 cells

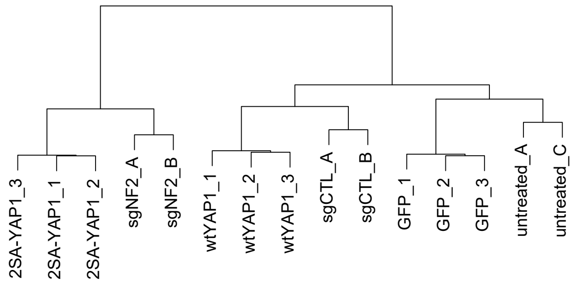


C Overlap between the DEGs of sgNF2 and 2SA-YAP1-expressing U5 human neural stem cells (compared to wild type cells)
FDR < 0.05, Fold Change > 1.5

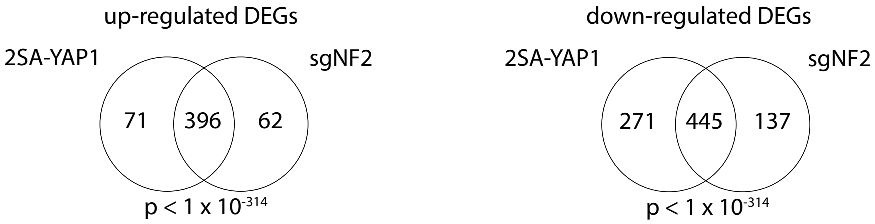


D Expression of YAP1 and YAP1 target genes in wild type (untreated), sgNF2, or 2SA-YAP1-expressing U5 human neural stem cells

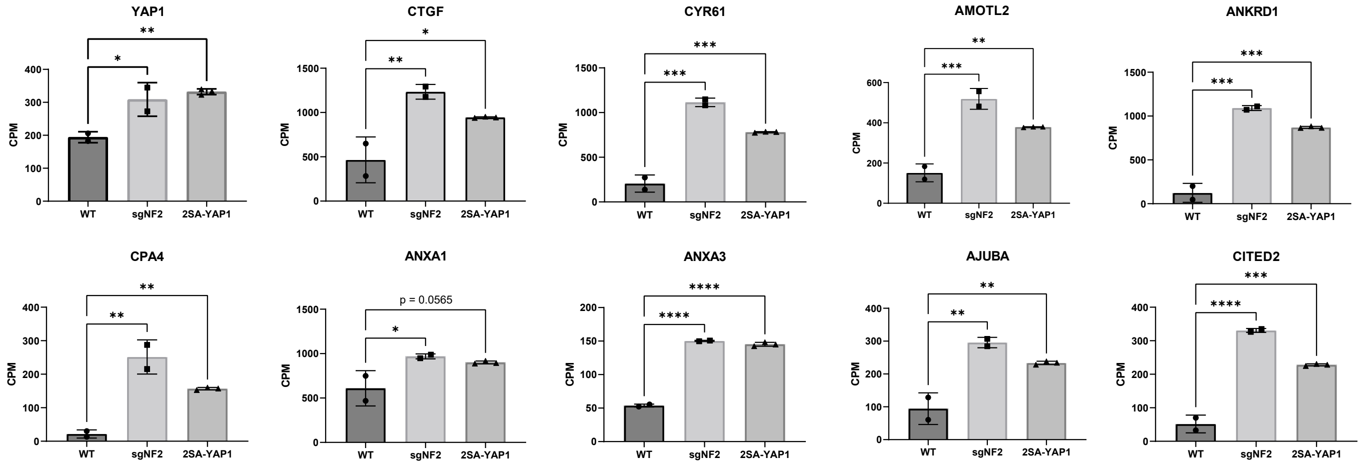

### Suppl. Figure S7E-U

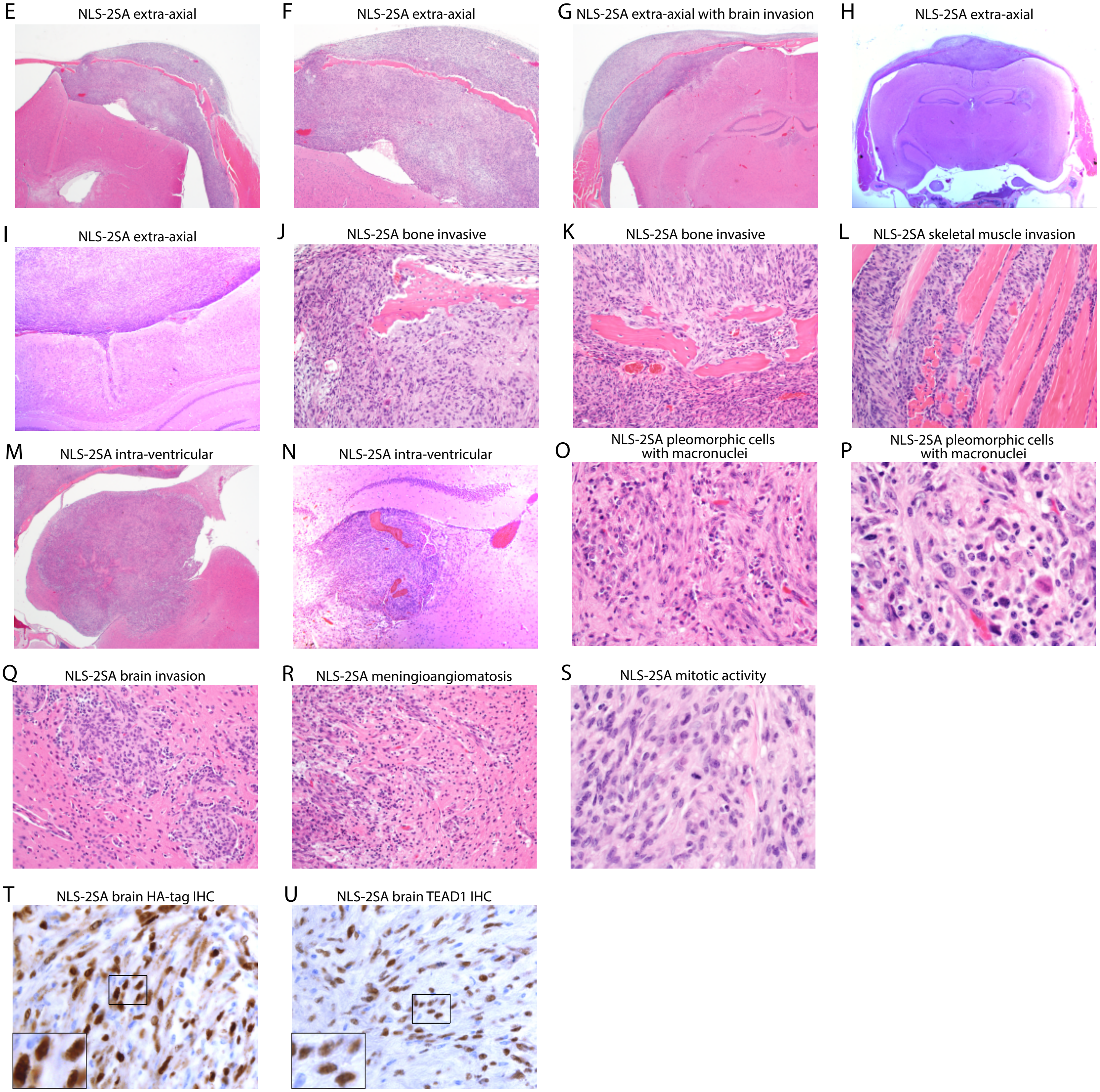

### Suppl. Figure S7V

#### V  UMAP Meningioma and PA samples - Calculated with complete dataset

UMAP Meningioma and PA samples - Dataset

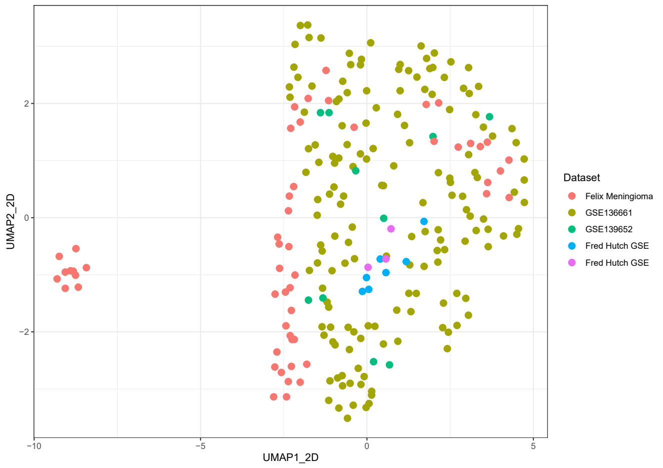


UMAP Meningioma and PA samples - NF2/Chr22 status

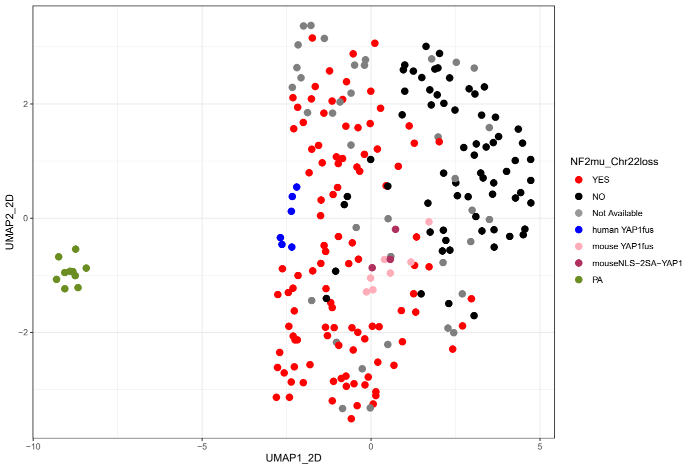


UMAP Meningioma and PA samples - Mutation status

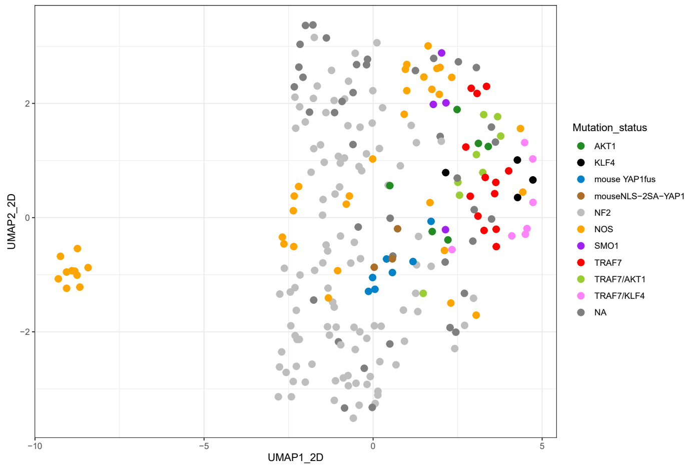


UMAP Meningioma and PA samples - Methylation subtype

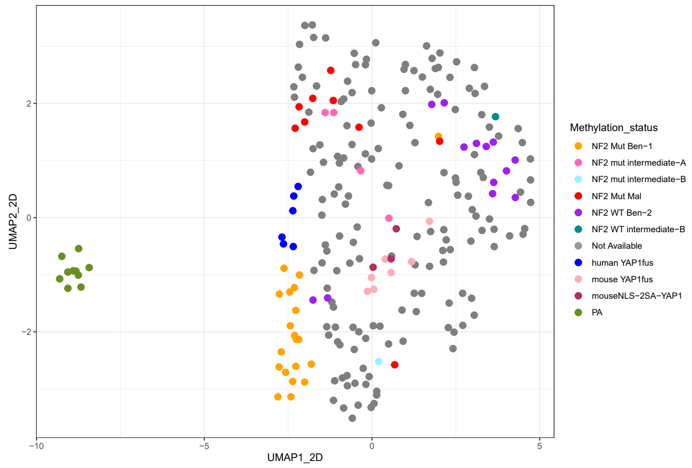

### Suppl. Figure S7W-Z

W Clustering of human, mouseYM, and mouse2SA samples based on expression of 2SA-YAP1 up-regulated genes

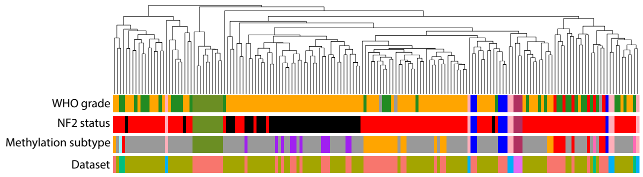


X Clustering of human, mouseYM, and mouse2SA samples based on expression of 2SA-YAP1 down-regulated genes

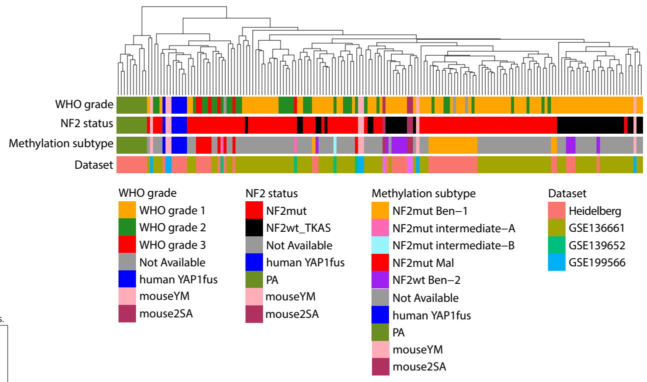


Y Expression of YAP1 target genes in human meningiomas, mouse YM tumors, mouse 2SA tumors, and PAs

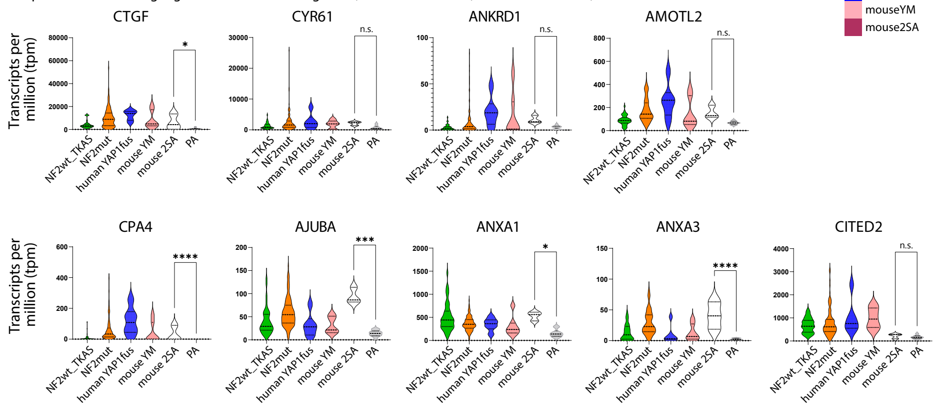


Z *ex vivo* treatment of NLS-2SA-YAP1-induced extra-cranial mouse tumors

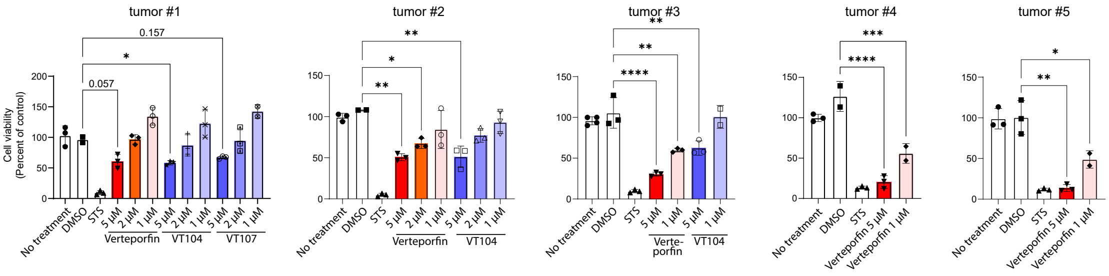

**Supplemental experimental procedures**

**Plasmid generation**

Primers used for plasmid generation are listed in Supplemental Table S3A. Additional plasmids used in this study are listed in Supplemental Table S3D.

**Transfection of RCAS viruses**

Chicken fibroblast (DF1) cells were maintained with 10% fetal bovine serum (FBS) in Dulbecco's modified Eagle medium (DMEM) (including 1% Penicillin/Streptomycin) at 39 degrees Celsius. DF-1 cells were transfected with the indicated RCAS plasmid using X-tremeGENE 9 DNA transfection reagent (Roche) according to the manufacturer's protocol. RCAS transgene expression was confirmed via western blot analysis.

**Magnetic resonance imaging (MRI)**

Tumor size and progression were evaluated using the preclinical 7 Tesla MR scanner (MR Solutions, DRYMAG7.0T) located in the FHCC imaging suite. Following anesthetic induction with 2% isoflurane, each mouse was placed on a bed and respiration was monitored via pneumatic pillow (SA Instruments, ERT gating module). Mouse body temperature was maintained by warm air circulation within the arm. Each procedure lasted approximately 10-30 minutes. Images were acquired using a mouse brain RF coil. Techniques used are Fast Spin Echo (FSE) T2 weighted scans (FSE T2w (axial) TE=75, 15 slices at 0.5mm thickness, FOV 25, 1 average) and for contrast enhanced imaging FSE T1 weighted scan pre- and post-contrast injection (FSE T1w (axial) TE=11, 15 slices at 0.5mm thickness, FOV 25, 2 averages). For contrast enhanced imaging 1 mL Gadobenate (MultiHance 529mg/ml, 0.5M) was diluted in 9 mL saline and mice were dosed with 4 ul/g of solution. Following imaging, each mouse was allowed to recover from anesthesia in a warm recovery cage.

**H&E Staining, Immunohistochemistry stainings of FFPE mouse tissues**

Mouse brains and hind limbs were formalin-fixed and paraffin-embedded, sectioned, and stained with H&E as described previously [6, 8]. Immunohistochemical staining of mouse brains and hind limbs was performed on the DISCOVERY XT platform (Ventana Medical Systems, Inc., Tucson, U.S.A) using the Discovery DAB Map Detection Kit according to standard protocols as described previously [7]. For a list of antibodies used see Suppl. Table S3B.

**Immunocytochemistry cellular protein localization**

HEK293 cells were seeded onto Laminin-coated 18 mm round cover slips (VWR International) at different cell densities, incubated overnight, and subsequently transfected with RCAS plasmids expressing HA-tagged constructs. 48 hours after transfection, cells were fixed with 2 percent para-formaldehyde, washed with cold PBS, permeabilized with 0.1 percent Triton-X-100, and subsequently stained for the HA tag (rabbit anti-HA, 1:2000, 3724, Cell signaling) as described previously [5]. For a list of antibodies used see Suppl. Table S3B.

**Luciferase assays**

HEK293 cells were cultured in DMEM, 10% FBS, 1% Penicillin/Streptomycin. If not indicated differently, HEK293 cells were seeded into white 96 well plates at 10,000 cells/well the day prior to transfection. Cells were then transfected with the indicated plasmids and a plasmid containing Renilla using Lipofectamin 3000 (Thermo Fisher Scientific) according to the manufacturer's instructions. Luciferase activity was measured 24 hours after transfection using the Dual-Glo Luciferase Assay System (Promega) on a Veritas Microplate Luminometer. For TEAD knockdown experiments, HEK293 cells were seeded into 96 well plates at 10,000 cells/well and incubated overnight and subsequently transfected with the indicated siRNAs (Suppl. Table S3C) on the next day. 24 hours later, the cells were transfected with the indicated RCAS plasmids, the GTIIC-Luc plasmid, and the Renilla plasmid. The Luciferase activity was measured 24 hours later as described above.

**Western Blot Analysis**

Cells were cultured, lysed, and processed for western blotting by standard methods. Proteins were resolved by SDS/PAGE (NuPAGE 10% Bis/Tris; LifeTech) according to XCell Sure Lock Mini-Cell guidelines, blocked with 5% milk/TBST and probed with specified antibodies overnight at 4°C in 5% BSA/TBST. After three TBST rinses, species-specific secondary antibodies were added in 5% milk/TBST. Blots were rinsed three times with TBST before being developed with Amersham ECL Western Blotting Detection Reagents (GE Healthcare). For a list of antibodies used see Suppl. Table S3B.

**Lentiviral production and infection**

For virus production, pLJM1 (Addgene) constructs containing the inserts of interest were transfected into 293T cells, along with psPAX and pMD2.G packaging plasmids (Addgene), using polyethylenimine (Polysciences). Fresh media was added 24 hours later and viral supernatant harvested 24 hours after that. For infection of NIH3T3 cells, $1 \times 10^5$ cells/well were seeded into 6-well plates. Lentivirus was used unconcentrated and cells were infected at a MOI<1 24 hours after seeding. 72 hours after seeding, selection was begun for cells successfully expressing the constructs using 1 µg/mL puromycin (for 3 days).

**Spheroid assay**

NIH3T3 cells (either untransduced, expressing wtYAP1, wtMAML2, or YAP1-MAML2) were seeded at $5 \times 10^3$ cells per well in a 96-well ultra-low attachment plates (Corning) in DMEM with 10% FBS and spun down at 1250 rpm for 10 minutes. After 4 days, cells were treated with DMSO or VP at indicated concentrations. Growth of spheroids was monitored using live cell imaging every 2 hours for 4 days in the Incucyte ZOOM system (Essen). Average phase object area ($mm^2$) was used for analysis.

**RNA-Seq**

For *in vivo* gene expression data from RCAS tumors, brain tumor tissues were dissected, flash frozen in liquid nitrogen, and subsequently crushed on dry ice. For *in vitro* gene expression data of transfected HEK cells, 30,000 cells per well were seeded in 48 well plates and transfected with 300 ng plasmid DNA 16 hours later. RNA was extracted 48 hours post-transfection.

RNA was extracted using the Qiagen RNeasy Mini Kit according to the manufacturer's instructions. Genomic DNA was removed by on-column DNase digestion. Total RNA integrity was checked using an Agilent 4200 TapeStation (Agilent Technologies, Inc., Santa Clara, CA) and quantified using a Trinean DropSense96 spectrophotometer (Caliper Life Sciences, Hopkinton, MA). RNA-seq libraries were prepared from total RNA using the TruSeq Stranded mRNA kit (Illumina, Inc., San Diego, CA, USA). Library size distribution was validated using an Agilent 4200 TapeStation (Agilent Technologies, Santa Clara, CA, USA). Additional library QC, blending of pooled indexed libraries, and cluster optimization was performed using Life Technologies' Invitrogen Qubit® 2.0 Fluorometer (Life Technologies-Invitrogen, Carlsbad, CA, USA). RNA-seq libraries were pooled (70-plex) and clustered onto an S1 flow cell. Sequencing was performed using an Illumina NovaSeq 6000 employing a paired-end, 50 base read length (PE50) sequencing strategy.

**RNA-Seq Analysis**

Aligning mouse RNASeq data

Raw sequencing reads were checked for quality using FastQC (https://www.bioinformatics.babraham.ac.uk/projects/fastqc/). RNA-seq reads were aligned to the UCSC mm10 assembly using STAR2 [3] and counted for gene associations against the UCSC genes database with HTSeq [1].

Obtaining and aligning publicly available datasets to hg38

Raw sequencing data for human meningioma samples were obtained from the data repository of the Dept. of Neuropathology at the University Hospital Heidelberg [10, 14], GSE139651 [12], and GSE136661 [11]. Raw sequencing data for human Pilocytic Astrocytoma (PA) samples was obtained from the data repository of the Dept. of Neuropathology at the University Hospital Heidelberg and is available upon request. Raw sequencing data for human neural stem cells were obtained from GSE116970 (WT (untreated), sgNF2, sgCTL) and GSE137040 (S127/397A-YAP1, GFP, wtYAP1). SRA files downloaded from GEO were converted to fastq files using fasterq-dump from the SRA tool kit. Raw sequencing reads were checked for quality using FastQC (https://www.bioinformatics.babraham.ac.uk/projects/fastqc/). RNA-seq reads were aligned to the UCSC hg38 assembly using STAR2 [3] and counted for gene associations against the UCSC genes database with HTSeq [1].

Batch correction and combining datasets
BiomaRt [4] was used to convert the mouse gene symbols to human orthologs. Protein coding genes present in both mouse and human datasets were used for further analysis. ComBat-seq [16] was used to correct batch effects when using multiple batches of RNASeq datasets.

RNASeq Expression Analysis
Log2 TPM counts were calculated on the adjusted RNASeq datasets, and used for constructing Uniform Manifold Approximation and Projections (UMAP) of the data in R. UMAPs were constructed using R package umap (https://cran.r-project.org/web/packages/umap/index.html). Heatmaps were made using R/Bioconductor package pheatmap (https://CRAN.R-project.org/package = pheatmap). Dendrograms were made in R using the package dendextend(https://cran.r-project.org/web/packages/dendextend/index.html). Volcano plots were made using ggplot2 in R [15]. Differential Expression analysis for RNASeq Data was performed using R/Bioconductor package DESeq2 [9] and edgeR [13]. A log2fold change cut-off of 0.58 (fold change of 50%) and FDR < 0.05 was used to find transcriptionally regulated genes. Gene Set Enrichment Analysis (GSEA) was performed against the MsigDB database with the GO Biological Process, GO Cellular Component, GO Molecular Function, KEGG, BioCarta, and Reactome gene sets using enrichR [2] package. Resulting enriched genesets and pathways were filtered via a threshold of FDR < 0.05. Fisher hypergeometric tests were implemented in R using function phyper() to see if genes in one set were over-represented, compared to other gene sets.

**Data and code availability**
RNA-Seq data can be accessed at NCBI Gene Expression Omnibus GSE199566. The code used to process and analyze the data is available at https://github.com/sonali-bioc/SzulzewskyYap1Maml2Paper. All other data associated with this study are present in supplementary materials and tables.

**Supplemental References**

1 Anders S, Pyl PT, Huber W (2015) HTSeq--a Python framework to work with high-throughput sequencing data. Bioinformatics 31: 166-169 Doi 10.1093/bioinformatics/btu638
2 Chen EY, Tan CM, Kou Y, Duan Q, Wang Z, Meirelles GV, Clark NR, Ma'ayan A (2013) Enrichr: interactive and collaborative HTML5 gene list enrichment analysis tool. BMC Bioinformatics 14: 128 Doi 10.1186/1471-2105-14-128
3 Dobin A, Davis CA, Schlesinger F, Drenkow J, Zaleski C, Jha S, Batut P, Chaisson M, Gingeras TR (2013) STAR: ultrafast universal RNA-seq aligner. Bioinformatics 29: 15-21 Doi 10.1093/bioinformatics/bts635
4 Durinck S, Spellman PT, Birney E, Huber W (2009) Mapping identifiers for the integration of genomic datasets with the R/Bioconductor package biomaRt. Nat Protoc 4: 1184-1191 Doi 10.1038/nprot.2009.97

5 Gurok U, Steinhoff C, Lipkowitz B, Ropers HH, Scharff C, Nuber UA (2004) Gene expression changes in the course of neural progenitor cell differentiation. J Neurosci 24: 5982-6002 Doi 10.1523/JNEUROSCI.0809-04.2004

6 Hambardzumyan D, Amankulor NM, Helmy KY, Becher OJ, Holland EC (2009) Modeling Adult Gliomas Using RCAS/t-va Technology. Transl Oncol 2: 89-95 Doi 10.1593/tlo.09100

7 Herting CJ, Chen Z, Pitter KL, Szulzewsky F, Kaffes I, Kaluzova M, Park JC, Cimino PJ, Brennan C, Wang Bet al (2017) Genetic driver mutations define the expression signature and microenvironmental composition of high-grade gliomas. Glia 65: 1914-1926 Doi 10.1002/glia.23203

8 Holland EC, Celestino J, Dai C, Schaefer L, Sawaya RE, Fuller GN (2000) Combined activation of Ras and Akt in neural progenitors induces glioblastoma formation in mice. Nat Genet 25: 55-57 Doi 10.1038/75596

9 Love MI, Huber W, Anders S (2014) Moderated estimation of fold change and dispersion for RNA-seq data with DESeq2. Genome Biol 15: 550 Doi 10.1186/s13059-014-0550-8

10 Maas SLN, Stichel D, Hielscher T, Sievers P, Berghoff AS, Schrimpf D, Sill M, Euskirchen P, Blume C, Patel Aet al (2021) Integrated Molecular-Morphologic Meningioma Classification: A Multicenter Retrospective Analysis, Retrospectively and Prospectively Validated. J Clin Oncol 39: 3839-3852 Doi 10.1200/JCO.21.00784

11 Patel AJ, Wan YW, Al-Ouran R, Revelli JP, Cardenas MF, Oneissi M, Xi L, Jalali A, Magnotti JF, Muzny DMet al (2019) Molecular profiling predicts meningioma recurrence and reveals loss of DREAM complex repression in aggressive tumors. Proc Natl Acad Sci U S A 116: 21715-21726 Doi 10.1073/pnas.1912858116

12 Prager BC, Vasudevan HN, Dixit D, Bernatchez JA, Wu Q, Wallace LC, Bhargava S, Lee D, King BH, Morton ARet al (2020) The Meningioma Enhancer Landscape Delineates Novel Subgroups and Drives Druggable Dependencies. Cancer Discov 10: 1722-1741 Doi 10.1158/2159-8290.CD-20-0160

13 Robinson MD, McCarthy DJ, Smyth GK (2010) edgeR: a Bioconductor package for differential expression analysis of digital gene expression data. Bioinformatics 26: 139-140 Doi 10.1093/bioinformatics/btp616

14 Sievers P, Chiang J, Schrimpf D, Stichel D, Paramasivam N, Sill M, Gayden T, Casalini B, Reuss DE, Dalton Jet al (2020) YAP1-fusions in pediatric NF2-wildtype meningioma. Acta Neuropathol 139: 215-218 Doi 10.1007/s00401-019-02095-9

15 Wickham H (2016) ggplot2: elegant graphics for data analysis. Springer New York, City

16 Zhang Y, Parmigiani G, Johnson WE (2020) ComBat-seq: batch effect adjustment for RNA-seq count data. NAR Genom Bioinform 2: lqaa078 Doi 10.1093/nargab/lqaa078
